# Nonequilibrium phases of a biomolecular condensate facilitated by enzyme activity

**DOI:** 10.1101/2024.08.11.607499

**Authors:** Sebastian Coupe, Nikta Fakhri

## Abstract

Biomolecular condensates represent a frontier in cellular organization, existing as dynamic materials driven out of equilibrium by active cellular processes. Here we explore active mechanisms of condensate regulation by examining the interplay between DEAD-box helicase activity and RNA base-pairing interactions within ribonucleoprotein condensates. We demonstrate how the ATP-dependent activity of DEAD-box helicases—a key class of enzymes in condensate regulation—acts as a nonequilibrium driver of condensate properties through the continuous remodeling of RNA interactions. By combining the LAF-1 DEAD-box helicase with a designer RNA hairpin concatemer, we unveil a complex landscape of dynamic behaviors, including time-dependent alterations in RNA partitioning, evolving condensate morphologies, and shifting condensate dynamics. Importantly, we reveal an antagonistic relationship between RNA secondary structure and helicase activity which promotes condensate homogeneity via a nonequilibrium steady state. By elucidating these nonequilibrium mechanisms, we gain a deeper understanding of cellular organization and expand the potential for active synthetic condensate systems.

## Introduction

Biomolecular condensates are dynamic intracellular structures that play a crucial role in organizing cellular biochemistry by locally concentrating proteins and ribonucleic acids (RNA) [1–3]. These condensates influence a range of cellular functions, from accelerating biochemical reactions to sequestering biomolecules and regulating polymer concentrations in the dilute phase [1–3]. While the principles of equilibrium polymer phase separation have been instrumental in describing many aspects of condensate formation and phase behavior, it is essential to recognize that these structures operate within a nonequilibrium cellular environment. As a result, biomolecular condensates are subject to kinetic trapping, aging effects [4–7], and can be driven out of equilibrium by chemically active processes, such as post-translational modifications, polymer production and degradation, and the active remodeling of intermolecular interactions [8–13]. This active nonequilibrium driving leads to novel behaviors in condensates, including size uniformity and resistance to Ostwald ripening [14, 15], asymmetric localization patterns [16], complex topologies [17, 18], dissolution [12, 13], and even droplet division and motion [15, 19]. A deeper understanding of the interplay between active cellular processes and the properties of biomolecular condensates is critical for manipulating these structures within cells and designing synthetic active condensates. Additionally, these active processes are fundamental to defining the evolutionary constraints and strategies that govern biomolecular condensation in cells.

DEAD-box helicases are a class of enzymes whose activity is central to biomolecular condensate regulation and homeostasis [13, 20]. The canonical biochemical role of DEAD-box helicases is in RNA processing, where they remodel RNA secondary structures and intermolecular interactions [21, 22]. DEAD-box helicases are non-processive RNA helicases that locally destabilize RNA base-pairing interactions, using adenosine triphosphate (ATP) hydrolysis to facilitate their cycling on RNA [21, 22]. More recently, these enzymes have gained interest due to their association with biomolecular condensates and the role their activity plays in facilitating normal condensate localization and structure in cells [12, 13, 20, 23, 24]. Their activity has also been tied to condensate dissolution and dynamics [12, 13, 20, 25, 26]. The mechanistic roles of this class of proteins in dictating condensate properties is an open area of interest and the capability of DEAD-box helicase activity to produce novel condensate behaviors is only starting to be realized.

Many biomolecular condensates are enriched in RNA and rely on RNA for their formation [27]. While RNA can fluidize biomolecular condensates in some contexts [28–30], RNA base-pairing interactions can also drive condensate formation, with extensive base-pairing shown to decrease condensate dynamics [31, 32]. Additionally, RNA secondary structures can influence both RNA incorporation into biomolecular condensates and the dynamics of the condensates themselves [33, 34]. This suggests that condensates containing RNA base-pairing interactions can be sustained out of equilibrium when coupled with energy-dependent helicase activity, which modifies RNA secondary structures and alters RNA-RNA interaction strengths.

To investigate the interplay between RNA-RNA interactions, DEAD-box helicase activity, and condensate properties, we engineered a reconstituted system composed of the LAF-1 DEAD-box helicase and a simple, repetitive, base-pairing RNA—a modified MS2 hairpin concatemer. Our results demonstrate that this system exhibits complex RNA concentration dynamics and gradients that are dependent on LAF-1 helicase activity. Increased helicase activity enhances RNA mobility within the condensed phase and prevents the formation of an RNA network. These findings establish a direct link between helicase activity and the metastable composition and dynamical state of condensates. Furthermore, disrupting defined RNA secondary structures induces a second RNA phase transition at the center of the LAF-1 condensed phase. This highlights how ATP-dependent enzymatic activity and RNA secondary structure counterbalance each other to maintain the system in a nonequilibrium steady state. Our work broadens the design principles of active synthetic and biological condensate systems and underscores the pivotal role of DEAD-box helicase activity as a nonequilibrium driver of ribonucleoprotein condensate properties.

## Results

### A model condensate system containing base-pairing RNA and a DEAD-box helicase exhibits time- and activity-dependent RNA gradients

To examine the interplay between DEAD-box helicase activity and RNA-RNA interactions in defining condensate properties, we generated a model hairpin multimer, based on a design used previously to study protein-RNA condensates [26], and combined it with a DEAD-box helicase known to form condensates in vitro, LAF-1 (Figure 1a) [25, 28]. These two components form protein-RNA co-condensates in the presence of ATP which are initially homogeneous (Figure 1b). However, over time, RNA gradients appear within the condensed phase, with higher RNA intensity at the center of the droplets (Figure 1 b,c). RNA fluorescence intensity decreases in the condensed phase over time suggesting RNA leaves the condensed phase (Figure 1 b,c). These RNA gradients emerge faster in smaller droplets, with smaller condensates also exhibiting a faster RNA decay in RNA fluorescence (Figure S1). Protein intensity remains homogeneous within the condensed phase and slightly increases in concentration over time (Figure 1b, Figure S1). We measured the RNA fluorescence decrease at the center of each condensate as a function of time (Figure 1d). Smaller droplets experience a faster decrease in RNA fluorescence intensity while larger droplets experience a slower decrease in RNA fluorescence intensity. The presence of gradients that are steeper at the boundary and condensate size-dependent fluorescence decay is consistent with RNA leaving from the droplet surface. This suggests that the process depends on the surface area to volume ratio of the condensate and should be renormalizable by a factor of droplet radius. When we renormalize the fluorescence timecourses by the radius of each condensate, we can collapse the fluorescence timecourse curves (Figure 1e). The decay portion of the collapsed fluorescence curve appears exponential in time, and we fit this curve to a single-exponential decay model (Figure 1e,f). The exponential fit timescales are similar across trials in the presence of ATP (Figure 1f). These results show that RNA leaves the condensate over time with a characteristic timescale that depends on a balance between surface area and the amount of RNA in a condensate.

**Figure 1:**
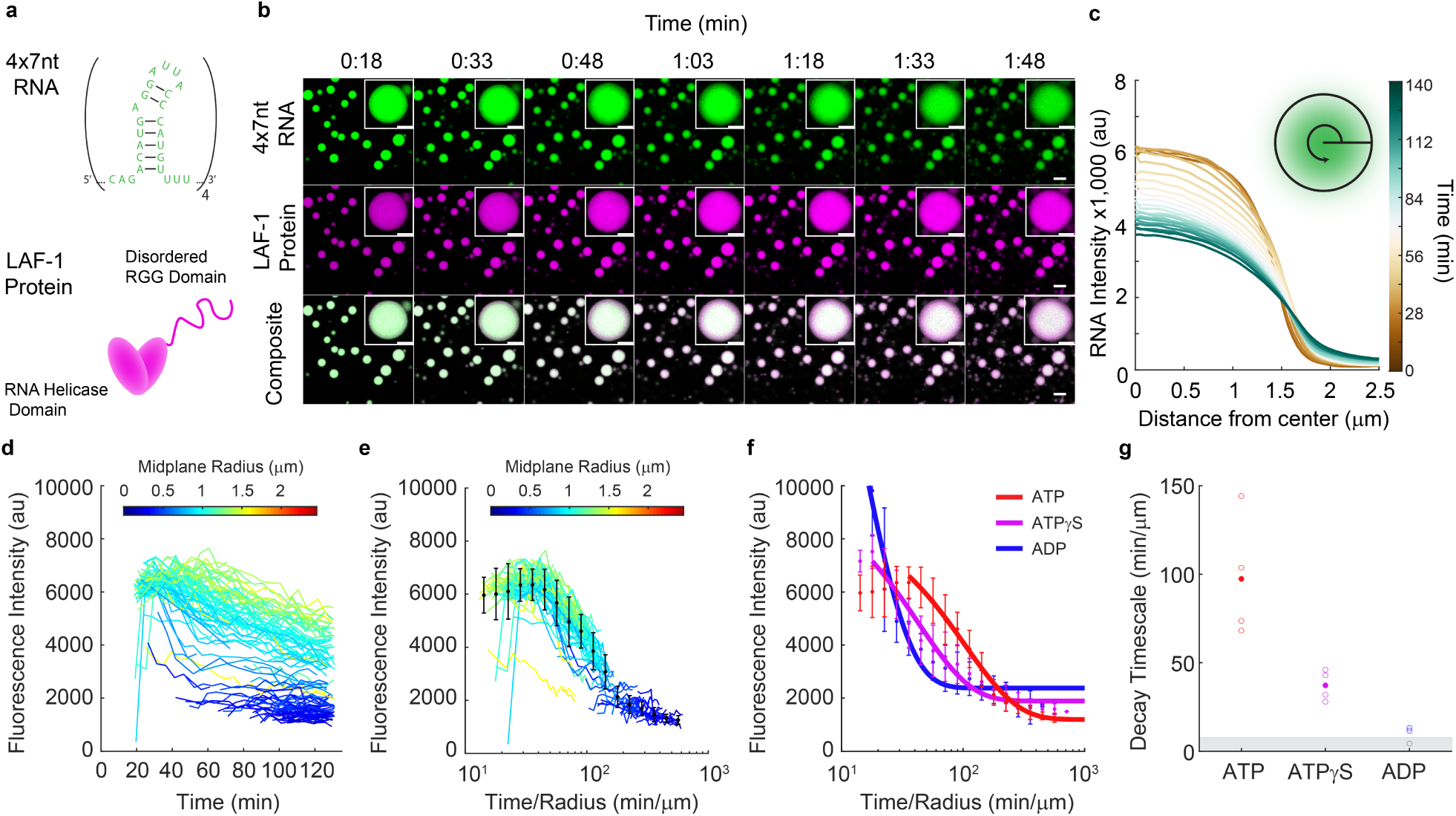
A ribonucleoprotein condensate exhibits time- and activity-dependent RNA gradients. a) Ribonucleoprotein condensates were formed from a MS2-hairpin 4x concatemer (top) and the LAF-1 DEAD-box RNA helicase (bottom) in the presence of ATP. b) LAF-1-4xMS2 RNA co-condensates formed in the presence of ATP are initially homogeneous, but over time RNA gradients develop and RNA fluorescence intensity decreases within the condensed phase. Top) RNA in the condensed phase produces radial gradients over time that depend on condensate size. Middle) Protein signal within the condensed phase is homogeneous and accumulates in the condensed phase over time. Bottom) Overlay of the channels shows radial-dependent RNA:LAF-1 ratios, with higher RNA to protein ratios at the center of the co-condensates. Scale bar = 5 *µ*m. The inset depicts these same phenomena for a single condensate, scale bar = 2 *µ*m. c) Average radial fluorescence intensity profiles for circular droplets with a mean radius of 1.956*µ*m +/- 0.122 *µ*m. 11 Droplets were included in the average. d) Average RNA fluorescence at the center of condensates with different sizes was tracked over time, with each trace color coded by the radius of the condensate’s midplane. All condensates experience a decrease in RNA concentration over time, though smaller condensates lose RNA faster, consistent with departure of RNA from the condensate surface. e) Renormalizing time by each condensate’s radius collapses the RNA fluorescence decay curves, consistent with a surface-area-to-volume-ratio dependent process. f) Renormalized fluorescence curves for LAF-1-4xMS2 RNA condensates formed in the presence of 1.6 mM ATP, ATP*γ*S, and ADP, with an exponential fit to the decay portion of the curve shown in solid lines. As helicase activity is impeded (ATP*γ*S) or abrogated (ADP), RNA departs more rapidly from the condensed phase. g) Fluorescence decay timescales extracted from exponential fits to the time-renormalized fluorescence decay curves. A slower departure of RNA from the condensed phase is seen with higher LAF-1 activity. Open circles represent fit timescales obtained for a single timecourse, filled circles are the mean of that condition. Decay timescales below around 10 min/*µ*m could not be calculated as the fluorescence had decayed to near the plateau value by the onset of the experiment and are represented as open circles within the grey box.

We next investigated ways to influence LAF-1’s helicase activity in order to use these perturbations to study the influence of activity in our reconstituted system. We avoided varying nucleotide concentration as this can have effects on condensate phase behavior through ATP’s chemical properties, independent of its identity as a substrate for LAF-1 [35]. Instead, we varied nucleotide substrate identity, which has a less pronounced effect than nucleotide concentration on biomolecular condensation [35]. To understand how nucleotide identify affects LAF-1’s helicase activity, we turned to a bulk fluorescence unwinding assay to study LAF-1’s unwinding of a short RNA duplex. The assay monitors the fluorescence of a fluorophore-quencher pair assembled on different strands of an RNA duplex (Figure S2a). We found that LAF-1 is capable of unwinding short RNA duplexes in the presence of ATP (Figure S2b). The ATP analog ATPγS slows helicase activity and also decreases the total amount of unwound RNA, while LAF-1 is incapable of unwinding the RNA substrate in the presence of ADP (Figure S2b, Table S1). We note that the finding that LAF-1 is capable of unwinding short stretches of duplex RNA contradicts the conclusion of a previously published study [36]. However, it is known that DEAD-box helicase activity is sequence independent but typically inversely related to the strength of a duplex and limited to short stretches of RNA [21, 22]. Therefore, we propose that the previous finding of LAF-1’s inability to unwind RNA was a product of the studies’ test substrate (18 base-pairs, 70% GC content) being too strong for LAF-1 [36]. The test substrate used in this study has a melting temperature that is higher than the predicted base-pairing energy of the MS2 hairpin (24°C vs 22°C) [37]. From these results, we conclude that LAF-1 can unwind sufficiently short duplexes of RNA and that this unwinding activity is related to its ATPase activity.

We next wanted to assess how the impeded LAF-1 helicase activity would influence the time-evolution of the RNA gradients we observe in the LAF-1:4xMS2 condensates. By impairing LAF-1’s helicase activity with ATPγS and ADP, we were able to generate faster time-evolution of the RNA gradients and faster fluorescence decay timescales within the condensed phase (Figure 1f, Figure S3). The fit timescale of fluorescence decay also becomes faster as helicase activity is impeded through ATP analogs or ADP. (Figure 1g). This is not due to a change in binding affinity for RNA (Figure S4, Table S2). Overall, these results demonstrate that RNA partitioning and homogeneity within the condensed phase is tied to LAF-1’s helicase activity.

Proper LAF-1 activity promotes RNA incorporation of the 4xMS2 hairpin and prevents the departure of RNA from the condensed phase.

### Preservation of RNA homogeneity correlates with preservation of RNA dynamics within the condensed phase

Though we found higher helicase activity corresponded to slower RNA departure from the condensed phase, we did not know whether helicase activity was slowing RNA diffusion to the condensate boundary or instead preventing RNA departure from the condensed phase. To test if the dynamics of the condensed phase were different as a function of unwinding activity, we performed fluorescence recovery after photobleaching (FRAP) on labelled LAF-1 protein and 4xMS2 RNA in the presence of ATP, ATPγS, and ADP. We performed the FRAP experiments on different condensates at different points in time to construct time-resolved estimates of biomolecular diffusion as a function of LAF-1 helicase activity (Figure 2a-c). RNA recovery dynamics within the condensed phase are initially faster and have similar rates in the presence of ATP, ATPγS, and ADP (Figure 2). However, over time, the RNA dynamics within the condensed phase slow down or age (Figure 2). This aging becomes pronounced for LAF-1:4xMS2 RNA condensates formed in the presence of ATP after 60-70 minutes (Figure 2a,d), while in the case of ATPγS occurs after only 40 minutes (Figure 2b,e), and in the case of ADP occurs after only 25-30 minutes (Figure 2c,f). Additionally, the recovery amplitude varies with helicase activity, with higher helicase activity preserving RNA recovery amplitude within the condensed phase for longer (Figure S5). The protein dynamics were constant over two hours with respect to both timescale and mobile fraction and were insensitive to unwinding activity (Figure S6). RNA dynamics are therefore tied to higher LAF-1 helicase activity while LAF-1 protein dynamics are not.

**Figure 2:**
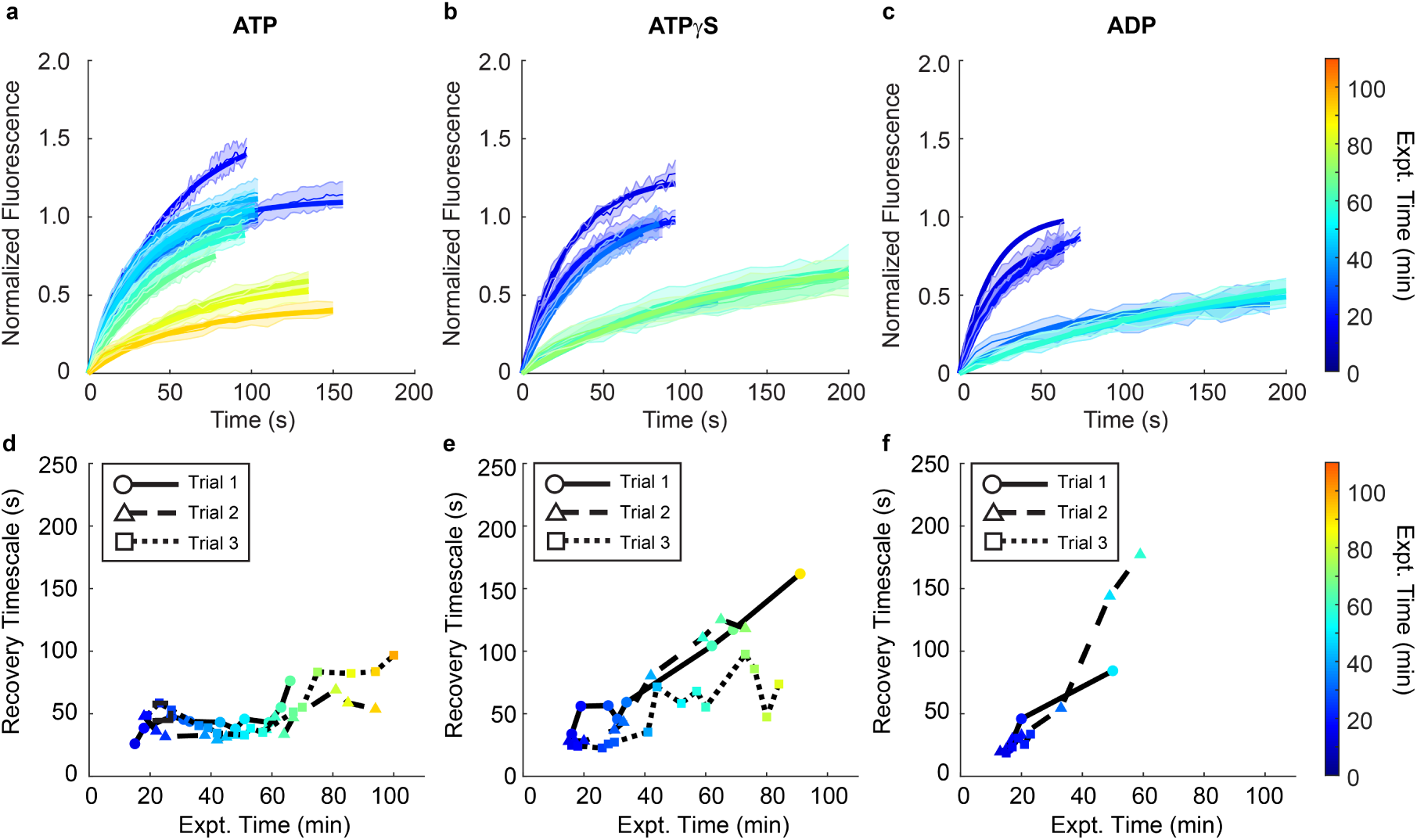
Hairpin RNA mobility within LAF-1 condensates decreases over time and depends on helicase activity. a-c) FRAP timecourse series of 4xMS2 hairpin RNA in LAF-1:RNA condensates formed in the presence of 1.6mM (a) ATP, (b) ATP*γ*S, or (c) ADP. Condensates formed in the presence of the different nucleotides have similar initial RNA recovery dynamics. However, a faster slowdown of RNA dynamics is seen as helicase activity is decreased. d-f) Recovery timescale as a function of the time at which the FRAP experiment was performed after system initialization for LAF-1:RNA condensates formed in the presence of (d) ATP, (e) ATP*γ*S, or (f) ADP. Data was compiled across three independent realizations. An increase in recovery timescale is seen for condensates formed with ADP and ATP*γ*S after 30-45 minutes of experiment time, with RNA dynamics slowest in the presence of ADP. Recovery amplitude fit parameters are shown in Figure S5.

That the RNA dynamics within the condensed phase were faster with higher helicase activity suggested to us that the longer RNA departure timescale from the condensed phase with ATP was due to a retention of RNA instead of slower RNA diffusion. Coupled to the faster unwinding activity and larger extent of unwinding with ATP seen in our helicase activity assays, this indicates that the formation of RNA secondary structures may promote RNA departure from the condensed phase. Consistent with this view, if we mix the system such that RNA can form hairpins before binding with LAF-1 protein, the RNA is excluded from the condensed phase and a pronounced RNA shell forms around LAF-1 droplets (Figure S7). This suggests that the timescale we extract in the fluorescence timecourses is a timescale related to the RNA folding rate in the system, which strongly depends on helicase activity. With higher LAF-1 helicase activity, there is a lower overall hairpin concentration at any point in time and thus a lower propensity for RNA to leave the LAF-1 condensed phase. With no or lessened LAF-1 helicase activity, hairpins form more quickly and leave the condensed phase.

### An RNA network forms within the condensed phase over time

At the end of the 2-hour timecourse, we noticed that the initially smooth RNA gradients had turned into punctate structures at the center of the condensed phase, and an RNA shell had emerged at the droplet periphery (Figure 3a-d). The brightest RNA punctate structures at the center of the condensed phase correlate with local minima in the LAF-1 protein signal, suggestive of an RNA network or phase that forms over the course of 2 hours (Figure 3e-h), Figure S8). These structures are visible both on a spinning disk confocal microscope and a Zeiss LSM980 laser scanning confocal with an AiryScan 2 detector (Figure S9). The AiryScan processing does not create the punctate structures as an artifact of processing strength (Figure S9). Importantly, these punctate structures do not form over time in the absence of the LAF-1 condensed phase, implying they form as a consequence of their high local concentration facilitated by the LAF-1 condensate (Figure S10). These observations are consistent with the recent finding that condensates containing high RNA concentrations can promote RNA aggregation through a percolation transition [38]. The formation of RNA punctate structures which exclude protein along with the slowed RNA dynamics over time suggest the formation of a percolated RNA network and a secondary RNA phase transition that competes with departure of RNA from the condensed phase.

**Figure 3:**
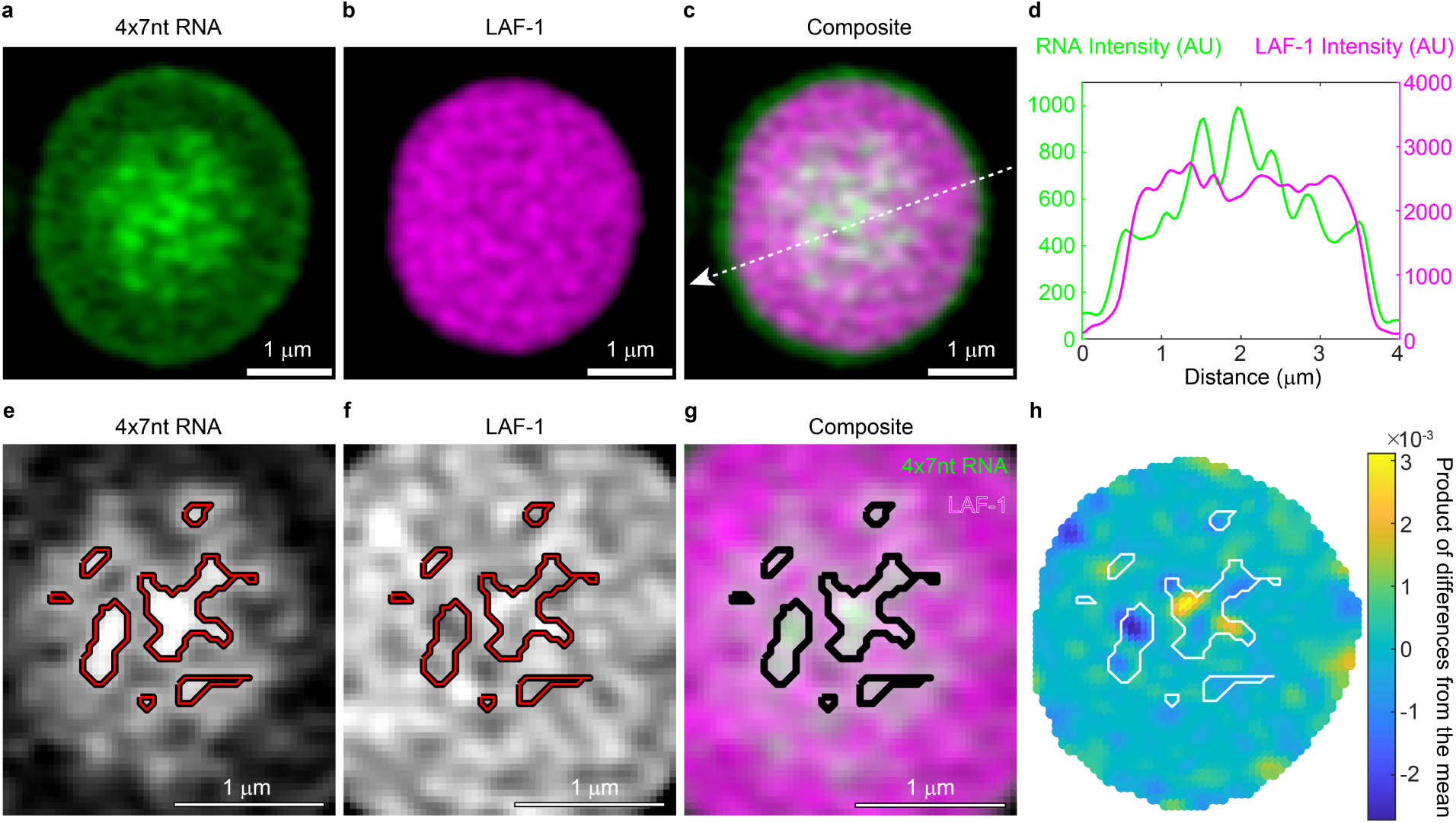
RNA punctate structures form over two hours and correlate with lower LAF-1 fluorescence intensity. a) 4xMS2 RNA fluorescence, b) LAF-1 protein fluorescence, and c) the composite image for a co-condensate that has aged for 2 hours. Bright RNA puncta are observed in the center of the droplet. Scale bar corresponds to 1 *µ*m. d) Line profile over the 4 micron slice shown in (c), which depicts fluorescence versus distance for LAF-1 protein and 4xMS2 RNA. LAF-1 and RNA intensity appear anti-correlated along the line profile, with highest RNA intensity and lowest LAF-1 fluorescence intensity visible at the center of the condensed phase. e-g) Segmenting out the brightest RNA features, these bright RNA regions correlate with local minima in the protein signal. Scale bar corresponds to 1 *µ*m. h) Normalized product of differences from the mean between the RNA and LAF-1 channels reveal that segmented RNA puncta overlay with negative correlation between the two channels. This suggests these RNA rich regions may be locally excluding LAF-1. Refer to Figure S8 for more examples of this phenomenon.

### RNA secondary structure prevents a second phase transition of RNA

To explore the idea of a competing RNA phase transition at the center of the LAF-1-RNA condensate, we reasoned that preventing RNA from leaving the condensed phase due to hairpin formation would increase the prominence of this event. We split the MS2 hairpin sequence into two separate constructs (4xMS2A, 4xMS2B) which each contain a section of the MS2 base-pairing motif (Figure 4a). Though both 4xMS2A and 4xMS2B simultaneously incorporate into the LAF-1 condensed phase to form initially homogeneous protein-RNA condensates, over the course of 90 minutes these constructs undergo a phase transition to an entirely RNA core which excludes LAF-1 (Figure 4b). This process also seems to depend on condensate size, with no RNA core seen for small droplets (Figure 4b). A finer timecourse of this process shows RNA gradients which are initially similar to the complete MS2 hairpin case but without the RNA departure from the condensed phase (Figure 4c). The RNA concentration at the center of the condensed phase continues to increase until there is a complete phase transition wherein the protein is completely excluded (Figure 4c). This effect does not rely on the specific base-pairing between the MS2A and MS2B motifs (Figure S11), which suggests it is due to nonspecific, weak base-pairing interactions between the RNA. This then implies the presence of the MS2 hairpin in earlier experiments prevents some of these transient interactions, either by decreasing the number of possible interaction sites or decreasing the conformational entropy of the RNA molecules. The 4xMS2A and 4xMS2B constructs are not capable of forming condensates under identical assay conditions in the absence of LAF-1 protein (Figure S12), meaning that either the presence of the LAF-1 condensed phase or the presence of LAF-1 helicase activity, or both, are required to promote this secondary RNA phase transition.

**Figure 4:**
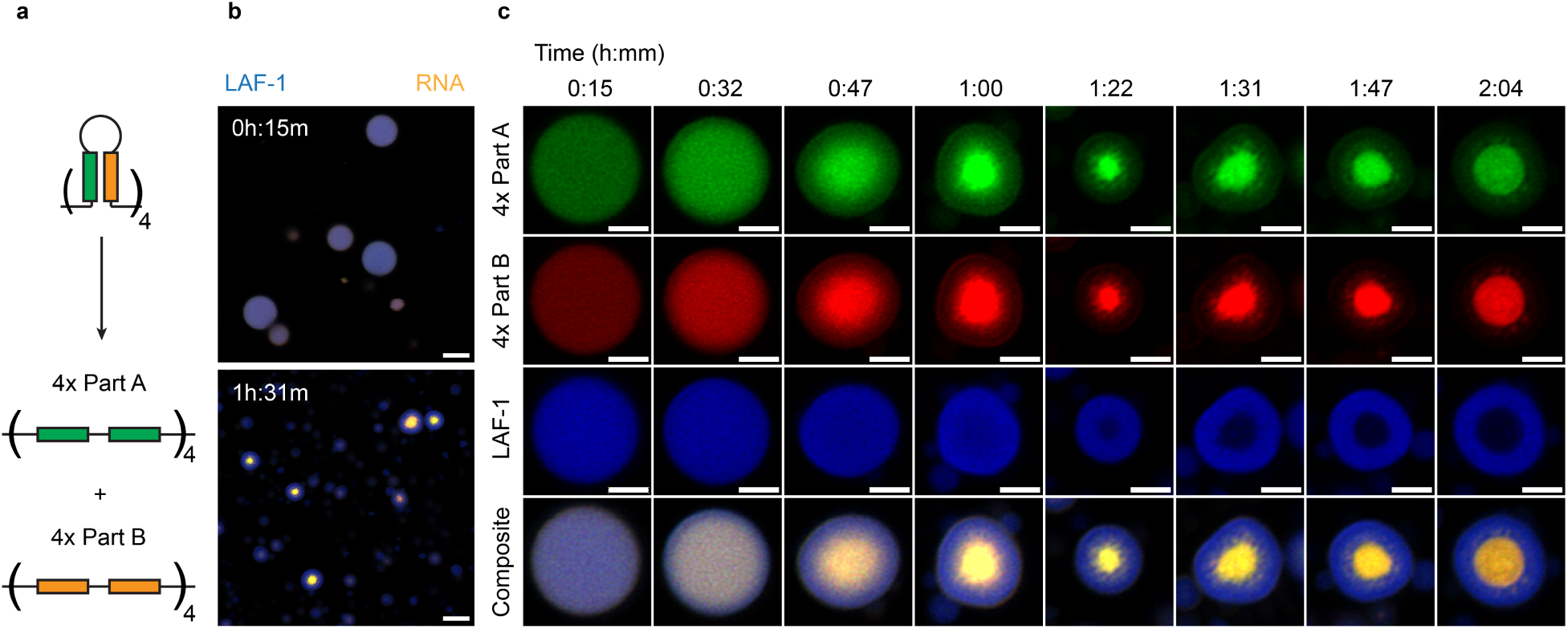
RNA constructs that lack stable RNA secondary structures undergo a second phase transition. a) The 4xMS2 hairpin construct was split into two separate constructs (Part A and Part B) that preserve the total base-pairing energy of the MS2 hairpin but which can only be satisfied intermolecularly. b) Condensates with both constructs are initially homogeneous in both protein and RNA (Top). After 90 minutes, the RNA in the system condenses at the center of the protein rich phase (Bottom). This second phase transition only occurs for condensates above a critical size. Scale bar corresponds to 5 *µ*m. c) Snapshots of single condensates of roughly 4 *µ*m diameter over two hours. Complete exclusion of LAF-1 protein from the RNA-rich phase occurs after 80-90 minutes. Scale bar corresponds to 2 *µ*m.

## Discussion

In this work, we have demonstrated that the antagonistic interplay between helicase activity and RNA secondary structure is crucial for preserving condensate homogeneity, dynamics, and composition. The presence of RNA hairpins facilitates the release of RNA from the condensed phase, while ATP-dependent helicase activity counteracts this by unwinding RNA secondary structures, thereby retaining RNA within the condensate (Figure 5). Concurrently, RNA hairpins inhibit multiple interactions among RNA molecules, preventing the formation of a separate RNA phase within the LAF-1 protein condensate (Figure 5). This dynamic tug-of-war ensures condensate homogeneity and promotes effective RNA incorporation, while also preventing the high RNA concentrations within the condensed phase from triggering a secondary phase transition.

**Figure 5:**
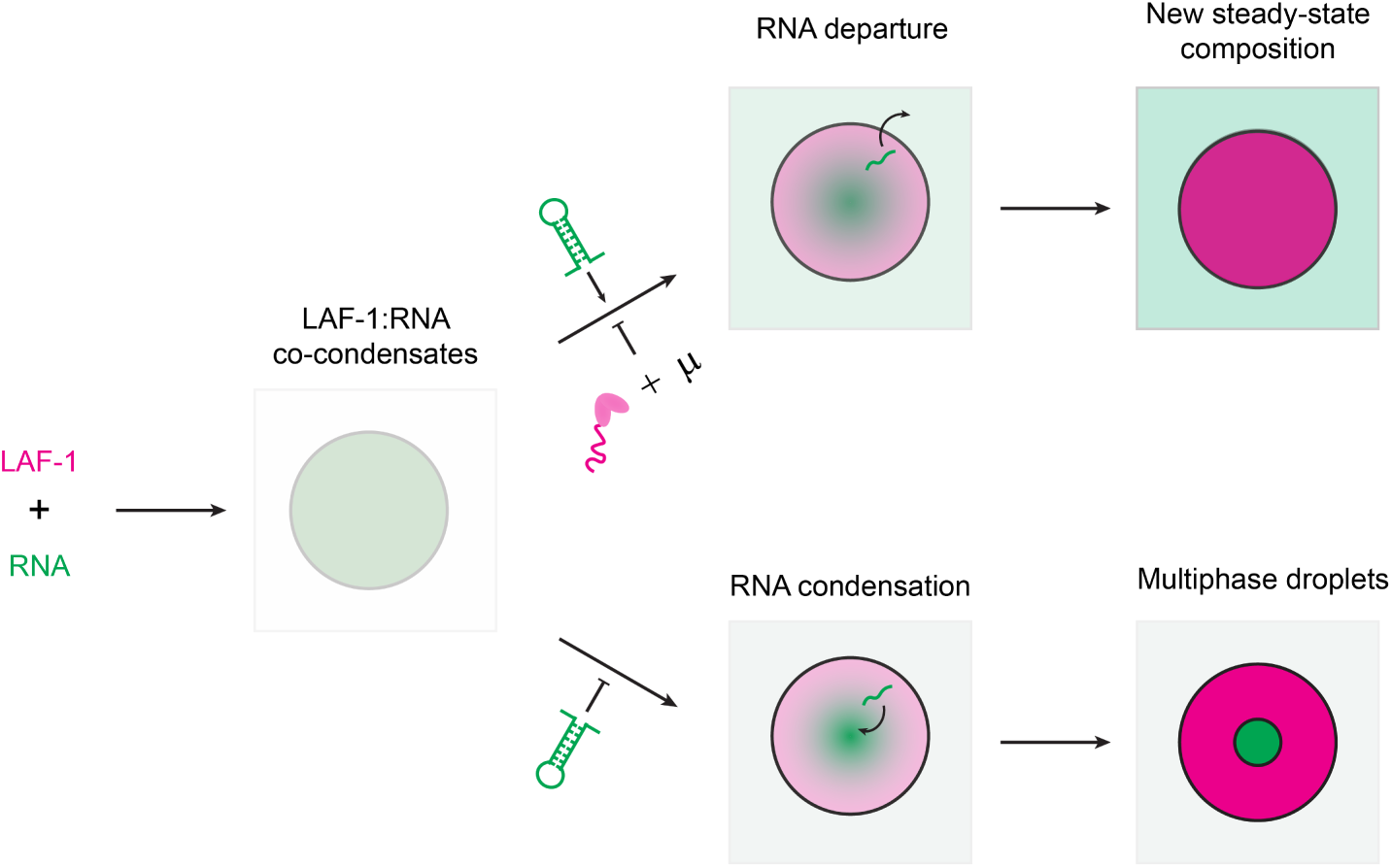
Model: Energy-dependent helicase activity and RNA secondary structure work in concert to preserve condensate homogeneity and dynamics. The formation of RNA secondary structure promotes RNA exclusion from the condensed phase. RNA helicase activity weakens or prevents RNA secondary structures from forming and therefore promotes RNA inclusion within the condensed phase. A condensation or aggregation of RNA occurs over time and is mitigated by the presence of RNA secondary structure, which decreases the number of potential RNA-RNA interaction sites, reduces RNA conformational entropy, or both.

These findings have significant implications for the biological role of DEAD-box helicases, particularly in promoting RNA incorporation into biomolecular condensates and preventing excessive local RNA concentrations from leading to aggregation or percolation transitions. Our results also suggest that RNA secondary structures may serve as an additional regulatory mechanism, allowing biological systems to favor RNA departure from the condensed phase when helicase activity alone is insufficient to prevent an RNA phase transition. Moreover, we show that the dynamics of biomolecular condensates containing base-pairing interactions can be actively maintained through energy-dependent helicase activity, which is likely critical for RNA processing and folding pathways localized to these cellular structures.

Finally, we have developed a novel biopolymer material whose internal enzymatic activity maintains its properties in a nonequilibrium steady state. By coupling enzyme activity to the interaction strengths that govern condensate formation and properties, we expand the design space for nonequilibrium condensates, transcending the sequence constraints of their constituent polymers. This work not only deepens our understanding of the fundamental principles governing cellular organization but also opens new avenues for the design of advanced synthetic condensate systems.

## Acknowledgements

We thank Sheena Vasquez for her feedback on the manuscript. We also thank Jaime Cheah and the Koch Institute’s Robert A. Swanson (1969) Biotechnology Center for technical support, specifically the High Throughput Sciences core, for training and assistance with the fluorescence polarization instrumentation. We would also like to thank Brandyn Braswell, Cassandra Rogers, and the W.M. Keck Microscopy Facility for access, training, and expertise in conducting the spinning disk and laser scanning confocal experiments. We thank the MIT Biology BioMicro Center for assisting with the capillary electrophoresis experiments. This work was supported in part by the Koch Institute Support (core) Grant P30-CA14051 from the National Cancer Institute, a graduate fellowship from MathWorks Inc., Sloan Foundation Grant G-2021-16758, and the Moore Foundation.

## Methods and Materials

### LAF-1 Expression, Purification, and Labelling

LAF-1 expression, purification and fluorescent labelling were performed as previously described [25, 28]. Briefly, the LAF-1 coding sequence was inserted into a pET28a(+) vector and was recombinantly expressed in *E. coli* BL21(DE3) cells following induction with 1mM IPTG. The BL21(DE3) cells were chemically lysed with lysozyme in the presence of Triton X100 and a Roche cOMPLETE protease inhibitor cocktail tablet, followed by mechanical lysis with a tip sonicator. The lysate was centrifuged and the soluble fraction retained. LAF-1 protein was purified using nickel affinity chromatography followed by heparin affinity chromatography. Purity and yield were assessed via SDS-PAGE and absorbance measurements at 280 nm. LAF-1 protein was then dialyzed into buffer containing 10% glycerol and flash frozen in liquid nitrogen.

To generate fluorescently labelled LAF-1 protein, Dylight633-NHS ester (Thermo Scientific 46417) was used following the manufacturer’s protocols. Protein was labelled prior to flash freezing.

### Unwinding Assays

Unwinding assays were designed and conducted as previously described [39]. RNA and DNA oligonucleotides were ordered from IDT as described in Table 1. Oligonucleotides were prepared as 100 µM stocks in nuclease free water.

**Table 1.**
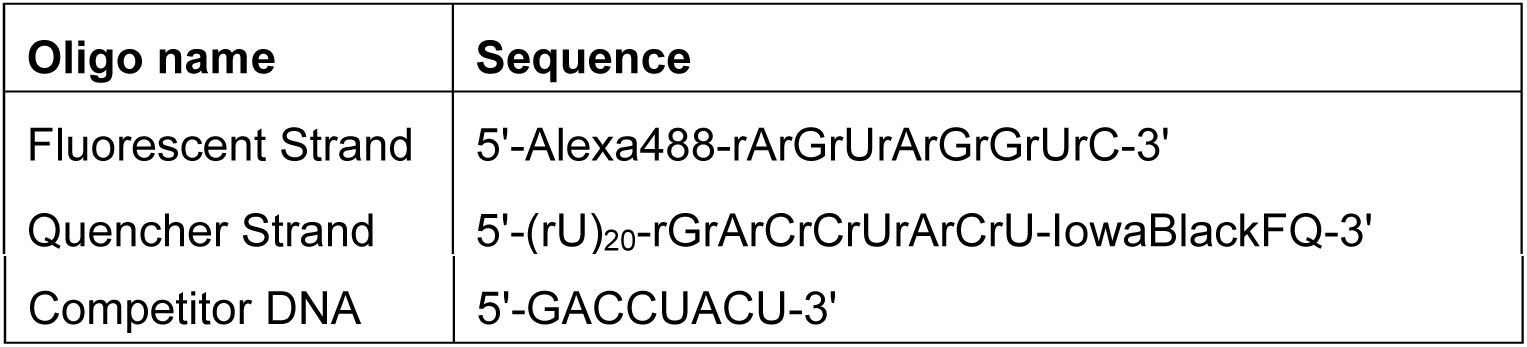
Sequences of oligonucleotides used in the unwinding assay.

RNA duplexes were prepared by annealing 200 µL of 1µM fluorescent RNA strand with 20 µL quencher RNA at 10 µM in 180 µL of reaction buffer (20 mM Tris pH 7.4, 2 mM magnesium acetate, 0.2 mM DTT). The reaction was heated to 80°C in a thermocycler and then cooled to 21°C in 3°C/3-minute increments. The reaction was spun down at 10,000xg for 10 seconds, then placed on ice for 5 minutes. 250 µL of the annealing mixture was then transferred to a fresh 1.7 mL microcentrifuge tube, along with 12.5 µL of 100 µM competitor DNA and 237.5 µL of 1x reaction buffer to produce a final duplex stock concentration of 250 nM.

LAF-1 protein was thawed at room temperature and buffer exchanged into fresh high salt buffer (20mM Tris pH 7.4, 1M NaCl, and 1 mM DTT). Protein 10x stock solution was then prepared at 5 µM in 20mM Tris pH 7.4, 1M NaCl, and 1mM DTT.

Fluorescence measurements were taken on a Tecan Infinite M200 Pro with fluorescence capabilities using an excitation filter of 488 nm with a bandwidth of 9 nm and an emission filter of 530nm with an emission bandwidth of 20nm. The gain was set at 125, 5 flashes were used for the measurement, each with an integration time of 1 msec.

A maximum intensity calibration sample was prepared consisting of 50 nM fluorescent RNA, 500 nM DNA competitor, 500 nM LAF-1, 2.5 mM ATP in reaction buffer. 70 µL of sample was transferred to a 96-well all-black fluorescent plate. Fluorescence readings were taken at 1 second intervals until a plateau in signal was reached, which was used as the maximum unwinding signal. Maximum calibration measurements were performed in triplicate.

Reaction mixtures were prepared with 50 nM annealed RNA and 500 nM LAF-1 in 1x reaction buffer. 63 µL of the mixture was transferred to the fluorescent plate and fluorescence readings were taken until a plateau was reached. At this point, 7 µL of 25 mM ATP was added to the mixture, the sample was mixed quickly, and then fluorescence readings were taken of the unwinding reaction. Reactions were performed in triplicate.

Fluorescence unwinding curves were normalized by first subtracting timecourse values from their reading at time = 0s. They were then divided by the maximum fluorescence calibration reading, less the intensity at time = 0s. Normalized curves were fit using the fit function in MATLAB. Two fit types were performed, a linear model of the initial rates and a single exponential fit. For the linear fit, an R^2^ value and RMSE were computed. For the exponential fit, 95% confidence intervals are reported for each fitting parameter (Table S1).

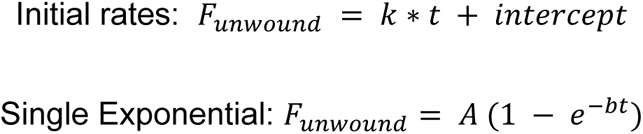

### Hairpin Construct Cloning

The pSL vector used for hairpin cloning was derived from the pSL-MS2-12x Addgene construct. pSL-MS2-12X was a gift from Robert Singer (Addgene plasmid # 27119 ; http://n2t.net/addgene:27119 ; RRID:Addgene_27119) [40].

The DNA sequences for single hairpins were ordered from IDT. They were then inserted into a pSL vector with a 5’ BamHI restriction site and a 3’ BglII restriction site. To create concatenated hairpins, we followed a procedure similar to the one used to originally clone the 12xMS2 hairpin construct [40]. A shorter concatemer number than 12x was desired as this significantly increased in vitro transcription yield while enabling study of base-pairing interactions. We first amplified the single hairpin using polymerase chain reaction (PCR), followed by restriction digest and gel purification. The DNA sequence for the single hairpin was then ligated with T4 ligase but in the presence of BamHI and BglII, in NEB buffer 3.1 with 1mM additional ATP. The BamHI and BglII sites have one nucleotide of overhang that is complementary. In this way, we achieved successful head-to-tail ligation which the restriction enzymes cannot cleave when the overhangs ligated in the BamHI-BglII site configuration. Reactions were allowed to proceed overnight at 37°C.

After overnight incubation, the samples were run on a 2% agarose gel and gel purified. PCR was then run on the samples to both increase concentration of the concatemers and introduce Gibson assembly overhangs on the concatemers. A second agarose gel and gel extraction step was performed on the products of the Gibson PCR. Many different concatemers were present at this step, and bands corresponding to the desired repeat number were selected. Gibson assembly was performed on the selected hairpin repeat constructs with the pSL vector backbone followed by a 4x dilution with water and transformation into NEB5α competent cells.

Single colonies were picked and grown in LB media before miniprepping and Sanger sequencing.

### In vitro transcription and RNA labelling

The DNA sequence for the desired RNA construct was amplified with PCR using a primer that also added a T7 promoter sequence in front of the hairpin sequence. PCR products were run on a 2% agarose gel and gel extracted using the Qiagen QIAQuick gel extraction kit.

In vitro transcription was performed using the NEB HiScribe T7 High Yield RNA synthesis kit. For fluorescently labelled RNA, 1 µL of either 10mM Cy3-UTP (APExBIO) or 10mM fluorescein-12-UTP (Roche) was included in the 20 µL in vitro transcription reaction. RNA was purified via lithium chloride precipitation. RNA stocks were diluted to 600 ng/µL with nuclease free water, flash frozen in liquid nitrogen, and stored at -80°C for no longer than a month before use.

### Condensate sample preparation

Unlabeled LAF-1 protein and LAF-1 labelled with Dylight-633 was thawed at room temperature before being buffer exchanged into fresh high salt buffer (20 mM Tris pH 7.4, 1 M NaCl, 1 mM DTT). Labelled LAF-1 and unlabeled LAF-1 samples were combined such that the final LAF-1 sample had less than or equal to 1% of the protein labelled. The LAF-1 sample was then diluted to 25µM in high salt buffer as a working stock for experiments. RNA samples were thawed and stored on ice until use.

Reactions were assembled by mixing 2 µL of RNA at 600 ng/µL with 1 µL of 3x-folding buffer containing 10 mM Tris pH 7.4 and 10 mM potassium acetate. RNA was heated to 95°C and then cooled to room temperature in 3°C/3 minute intervals. The 3 µL RNA sample was then mixed with 3.2 µL of LAF-1 protein at 25µM in 1M NaCl, 20 mM Tris pH 7.4, and 1 mM DTT. This mixture was quickly added to another mixture containing 7.8µL of 20 mM Tris pH 7.4, 1 mM DTT and 1.2µL of 25 mM Mg+nucleotide in 20 mM Tris pH 7.4. The final concentrations of components were then 5 µM LAF-1, 75 ng/µL of RNA, 1.6 mM Mg nucleotide, 20 mM Tris, 200 mM NaCl, and 1 mM DTT.

Samples were incubated for 8 minutes before transferring 10 µL to a PDMS chamber adhered to a 1.5H coverslip that had been passivated overnight with 2% w/v Pluronic F-127. The chamber was then mounted on either an Andor Spinning Disk confocal or a Zeiss LSM980 laser scanning confocal.

### Andor Spinning Disk Fluorescence Imaging

Spinning disk fluorescence imaging was performed on a Nikon Ti-E inverted microscope equipped with an Andor iXon 897E EM-CCD camera, a Yokugawa CSU-X1 Spinning Disk Confocal Scan Head. A 100x Nikon objective with a numerical aperture of 1.4 was used for all spinning disk imaging.

### Zeiss Airyscan 2 Fluorescence Imaging

Laser scanning confocal imaging was performed on a Zeiss LSM980 laser scanning confocal equipped with a 63x, 1.4 NA, oil objective and an Airyscan 2 detector. Airyscan processing was done with default processing settings. Image grids of 3×3 (Big Image) were used for imaging to increase droplet imaging throughput without producing long time-delays vertical slices. Imaging intervals of 5 minutes between Big Images were used for all datasets. Imaging was done at 21-22°C.

In analyzing the timecourses, individual condensates were segmented and tracked in MATLAB using elements of the Image Processing Toolbox. Information about condensate fluorescence and shape could then be extracted using custom MATLAB scripts.

Radial profiles were generated in 2-dimensions by averaging pixel intensities at rings with increasing radius from the identified condensate center. Radial profiles were constructed from condensates whose midplanes had an average circularity of at least 0.9 and averaged radial profiles were computed using a radial bin size of 0.24 μm.

To measure the intensity at the center of each condensate, the midplane of each condensate was identified by finding the z-slice with the maximum number of pixels. The centroid of this midplane was then calculated and the average pixel intensity within one pixel of the centroid was calculated. Intensity values for the same condensate at different points in time were linked via trajectory construction performed on the midplane centroids. Intensity as a function of time curves were fit to single exponential decay models using Nonlinear Least Squares Curve Fitting in MATLAB.

For colocalization analysis, droplets were first segmented relative to the protein channel. A mask was applied using the protein contour, followed by a second segmentation based on the RNA channel. This second segmentation procedure was used to identify RNA puncta in the condensed phase. Colocalization between RNA and protein signal within the droplet contour was calculated using a normalized product of differences from the mean:

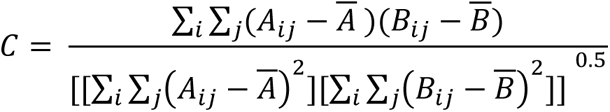

Where *i* and *j* denote row and column of the images, A denotes Channel 1 and B denotes Channel 2 of the image, and *A̅, B̅* are the averages of the Channel 1 and Channel 2 images, respectively.

### Fluorescence recovery after photobleaching

Fluorescence imaging was performed on a Nikon Ti-E inverted microscope equipped with an Andor iXon 897E EM-CCD camera, a Yokugawa CSU-X1 Spinning Disk Confocal Scan Head, and an Andor FRAPPA photomanipulation system. A 100x Nikon objective with a numerical aperture of 1.4 was used for all fluorescence imaging. For FRAP experiments, a spot size of 3 pixels, corresponding to roughly 0.42 µm, was used for photobleaching. FRAP timecourses were generated by FRAPping several droplets within a field of view to get statistics on recovery timescale at a particular time-point. Different fields of view were then imaged at successive time-points to generate information about recovery timescale as a function of the time at which the experiment was performed.

Individual droplets which had FRAP regions were cropped and bleach spots were identified using custom Python scripts (https://github.com/stcoupe/FRAP-analysis). Intensity within the bleached region was normalized relative to the intensity outside the bleach spot but within the droplet. We then performed an affine transformation of the data in order to be able to directly compare FRAP curves. The intensity ratio at the first bleach time-point (*I*(0)) was subtracted from each time-point (*I*(*t*)), and this was then divided by the difference between the average intensity ratio before bleaching (⟨ *I*_*i*_⟩) and *I*(0).

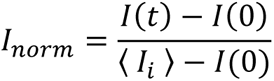

Note that this is a different normalization procedure than typical for condensate measurements [41]. However given the gradients present particularly in the RNA channels of our condensed phases, the modified normalization procedure was used.

Individual droplet FRAP curves were then fit by single exponentials using Nonlinear Least Squares fitting in MATLAB to measure the timescale and amplitude of fluorescence recovery.

### Fluorescence Polarization

An RNA oligo of rU20 with a 5’-Alexa488 fluorophore was synthesized by IDT.

LAF-1 protein was thawed at room temperature before being buffer exchanged into fresh high salt buffer (20 mM Tris pH 7.4, 1 M NaCl, 1 mM DTT). Protein samples were then diluted to 4 µM with high salt buffer and then concentration measurements were then taken in triplicate using a Tecan M200 plate reader with the Nanoquant insert. The samples were then diluted 5-fold with low salt buffer (20 mM Tris pH 7.4, 1 mM DTT) to produce the stock protein concentrations for the serial dilution.

Serial dilutions of 1.63x were prepared by filling the wells of a Corning 96-well flat bottom black plate with 105 µL of assay buffer (20 mM Tris pH 7.4, 200 mM NaCl, 1 mM DTT) containing 2 nM fluorescent RNA oligo and the desired concentration of nucleotide. 180 µL of protein stock solution containing 2 nM fluorescent RNA oligonucleotide and the desired concentration of nucleotide was then added to the first well, and successive volumes of 180 µL were transferred to subsequent wells. Samples were incubated for 5 minutes before fluorescence polarization readings were taken. Incubation time was tested to ensure the system had equilibrated by this point. Experiments were performed in triplicate.

Fluorescence polarization measurements were performed with a Tecan Infinite M1000Pro, using a 470nm excitation filter with a 5 nm bandwidth, a 520nm emission filter with a 10 nm bandwidth, 110 gain, and 10 flashes per reading. The G-factor for the fluorescence polarization was calibrated using fluorescein, and determined to be 1.162.

Fluorescence polarization as a function of protein concentration was plotted and fit to a Hill equation using Nonlinear Least Squares Fitting performed in MATLAB. A vertical offset (b) and amplitude term (A) were included to fit the data:

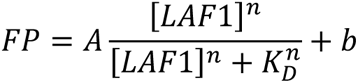

Fit parameters and their 95% confidence intervals can be found in Table S2.

## Supplementary Information

**Table 1:** Kinetic parameters for LAF-1 duplex unwinding assays performed with different nucleotides. Parameters were extracted from either a linear fit to the initial unwinding rates (columns 2-4) or single exponentials of the entire curve (columns 5 and 6). R^2^ values report on the goodness of fit for the linear fit parameters in columns 2 and 3. For the exponential fits, 95% confidence intervals of fit parameters are displayed in parentheses.

| Condition | $k_{initialrates}$<br>(fraction/s) | Intercept<br>(fraction) | Linear $R^2$ | Exp. A<br>(fraction) | $k_{exponential}$<br>(fraction/s) |
| --- | --- | --- | --- | --- | --- |
| ATP | 0.0134 | -0.0186 | 0.996 | 0.777 (0.763, 0.790) | 0.0246 (0.0232, 0.0259) |
| ATPγS | 0.00224 | -0.0162 | 0.983 | 0.289 (0.286, 0.291) | 0.0107 (0.0103, 0.0111) |
| ADP | -0.00023 | -0.00354 | 0.724 | -0.0179 (-0.0186, -0.0173) | 0.0354 (0.0239, 0.0470) |

**Table 2:** Apparent binding affinity parameters for LAF-1 fluorescence polarization assays in the presence of different nucleotides and their 95% confidence intervals. Fluorescence polarization curves in Figure S4 were fit to a Hill model where n was included as a fitting parameter except in the case of ATP*γ*S where it was fixed at 1.

| Condition | $K_{D, App}$ (nM) | $K_{D, App}$, 95% CI (nM) | n | n, 95% CI |
| --- | --- | --- | --- | --- |
| Buffer | 25.2 | (23.1, 27.2) | 1.68 | (1.48, 1.88) |
| MgCl<sub>2</sub> | 22.6 | (20.3, 24.8) | 1.95 | (1.62, 2.28) |
| MgATP | 11.2 | (9.44, 13.0) | 0.85 | (0.73, 0.96) |
| MgATPγS | 14.6 | (6.5, 22.7) | 1 | - |
| MgADP | 8.53 | (5.06, 12.0) | 0.54 | (0.39, 0.70) |

---

**Figure S1:**
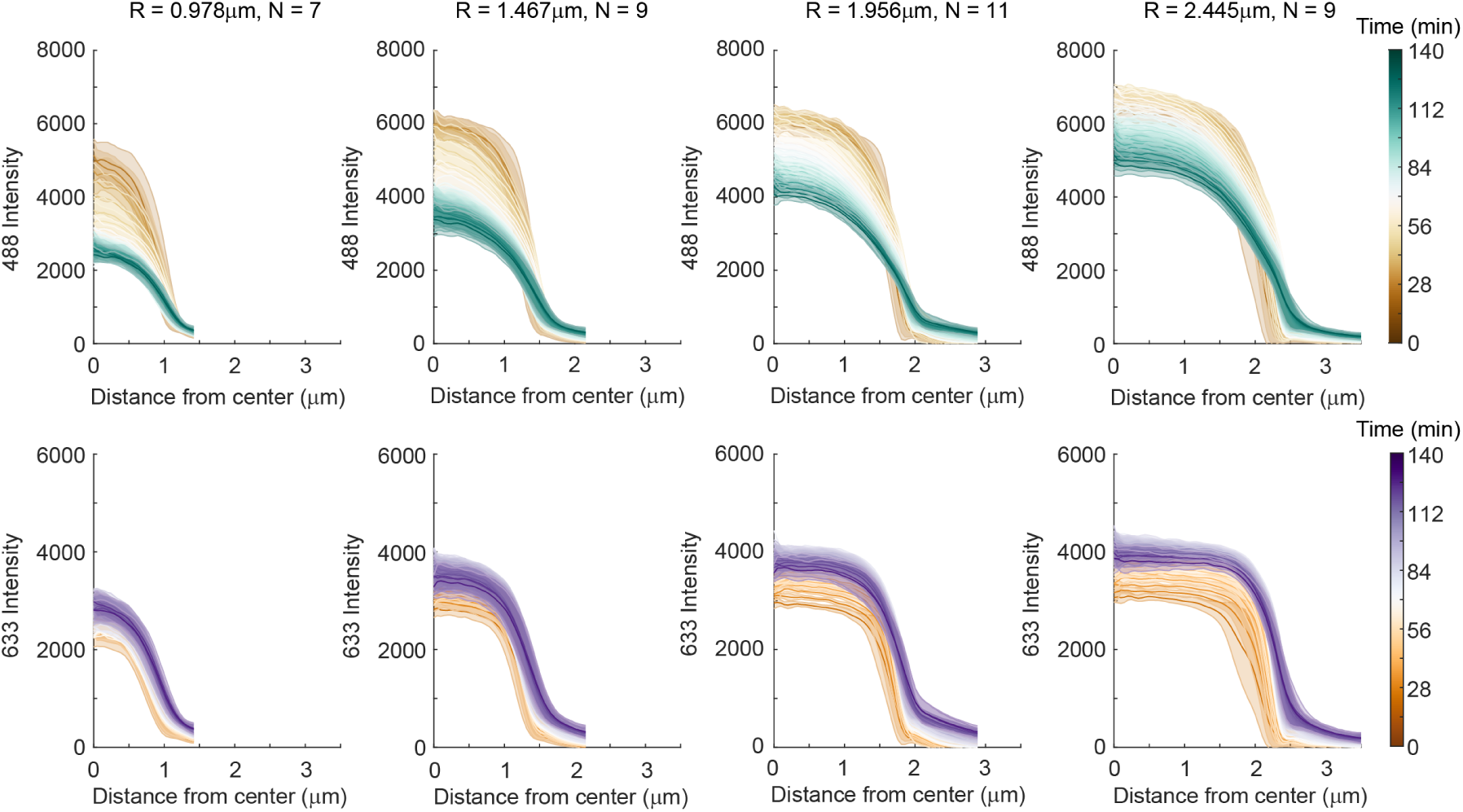
Average radial fluorescence intensity profiles for circular droplets of various sizes. Number of droplets included in the average and the radial bin center are indicated at the top. Each bin has a width of 0.245 *µ*m. Top Row) Average 4xMS2 RNA radial intensity profiles as a function of time. Gradients emerge more quickly and RNA fluorescence decreases faster in condensates of smaller sizes. This is consistent with RNA leaving from the condensate boundary. Bottom Row) LAF-1 radial intensity profile as a function of time. Protein profiles spatially match those of RNA at early times. No gradients are present in the LAF-1 profiles until the condensate boundary is encountered. Protein fluorescence increases over time.

**Figure S2:**
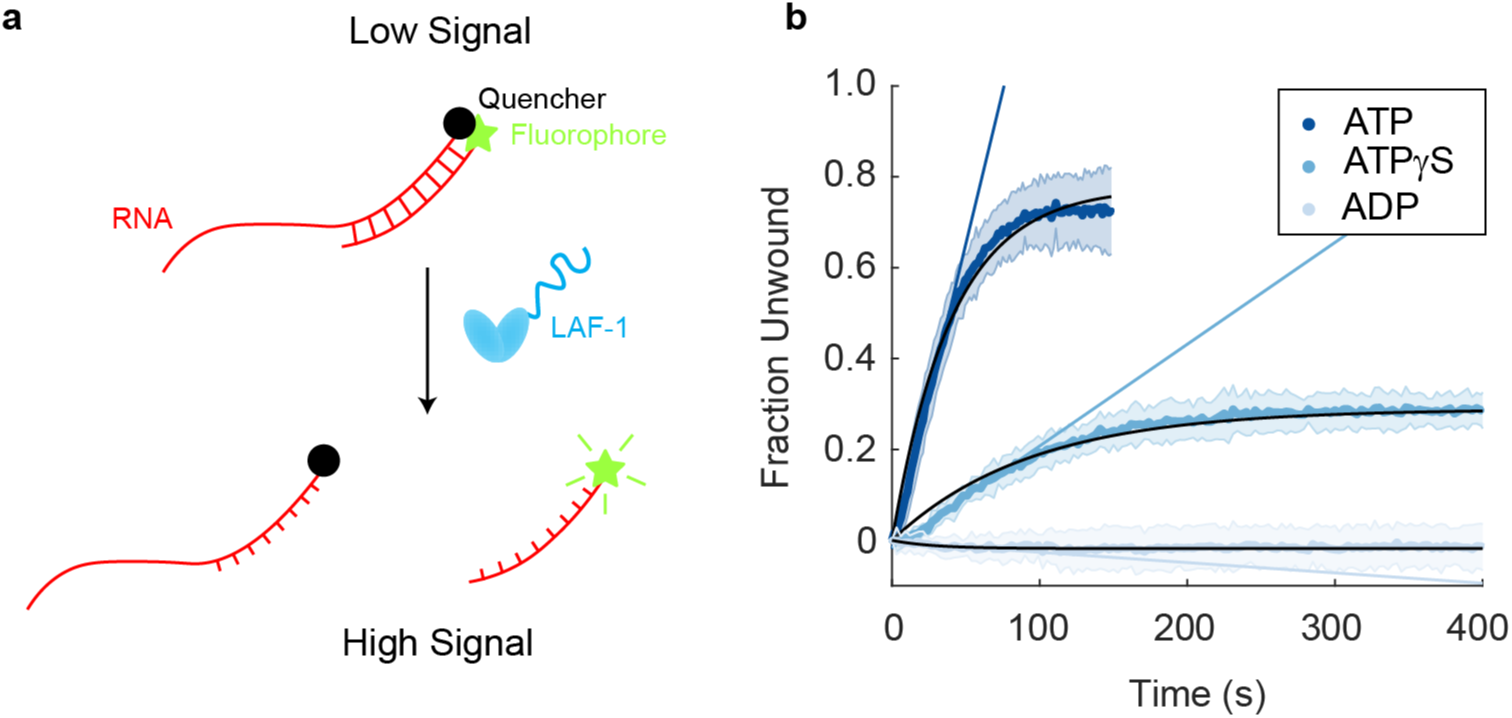
LAF-1 can unwind a short RNA duplex in an ATP-dependent manner. a) Overview of the real-time fluorescence-based unwinding assay [1]. Two RNA strands are annealed such that a fluorophore-quencher pair are in close proximity in the duplexed configuration. One of the RNAs contains a ssRNA overhang, known to be important for other DEAD-box helicases’ unwinding activities [2]. Upon unwinding, the fluorophore is liberated and the sample increases in fluorescence emission. b) Normalized unwinding curves from an unwinding assay performed on an RNA construct with a 50% GC content, 8-bp duplex and a U_20_ ssRNA overhang. Assays were performed in the presence of 500nM LAF-1, 100 mM NaCl, and 2.5mM of the indicated nucleotide. Fastest unwinding rate and highest unwinding fraction are seen with the system containing ATP. ATP*γ*S results in slower unwinding and a lower overall fraction of RNA unwound. ADP does not enable RNA unwinding by LAF-1. Fits were performed to single exponentials (solid black lines) and linear fits of the initial unwinding rates (solid colored lines) (Table S1).

**Figure S3:**
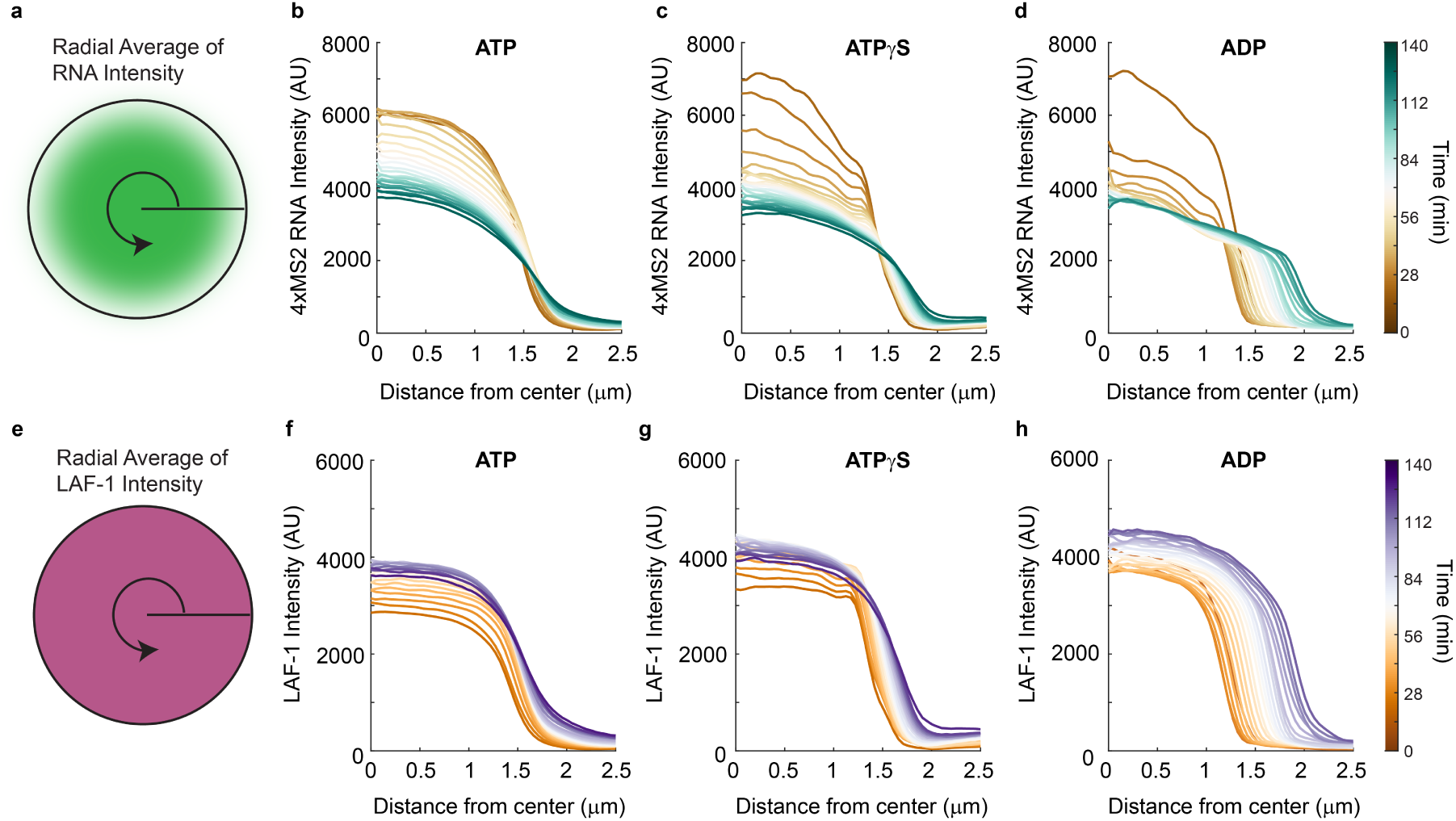
Radial gradient time-evolution depends on nucleotide identity. a) Average radial RNA intensity profile for LAF-1:4xMS2 RNA co-condensates formed in the presence of (b) ATP, (c) ATP*γ*S, or (d) ADP. A faster evolution of RNA gradients and RNA fluorescence decay are seen as helicase activity is impeded with ATP*γ*S and abrogated with ADP. e) Average radial LAF-1 protein intensity profile for LAF-1:4xMS2 RNA co-condensates formed in the presence of (f) ATP, (g) ATP*γ*S, or (h) ADP. Protein intensity increases in the condensed phase with ATP, ATP*γ*S, and ADP, with no spatial variation of protein signal within the condensed phase, and an increase in signal over time. Number of droplets included in each average: (b,f) 9 for ATP, (c,g) 9 for ATP*γ*S, (d,h) 6 for ADP. All droplets averaged had a radius of 1.71 *µ*m +/- 0.122 *µ*m and an average circularity of *>* 0.9.

**Figure S4:**
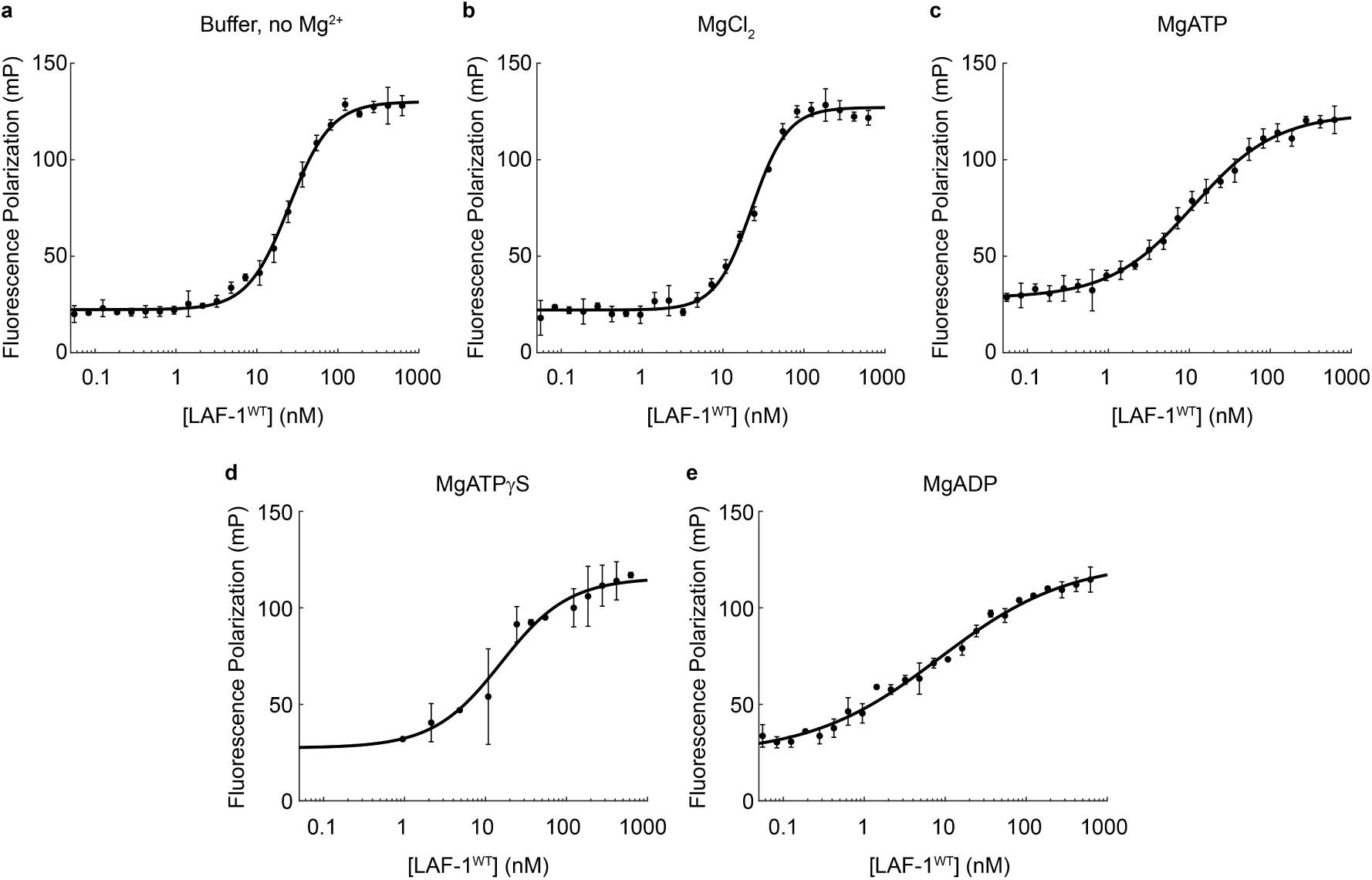
LAF-1 binds RNA with the same apparent affinity in the presence of a variety of nucleotides, but nucleotide presence decreases cooperativity of RNA binding. (a-e) Fluorescence polarization curves comparing LAF-1 binding to rU_20_ in (a) buffer (b) buffer + 1.6mM MgCl_2_, (c) 1.6 mM MgATP, (d) 1.6 mM MgATP*γ*S, or (e) 1.6 mM ADP. Error bars are the standard deviation over three independent replicates. Solid line represents the fit of the data to a Hill binding model, where cooperativity or n was included as a fit parameter, except for ATP*γ*S where n was fixed at 1. The apparent dissociation constants, Hill coefficients, and 95% confidence intervals for the fit parameters displayed in Table S2.

**Figure S5:**
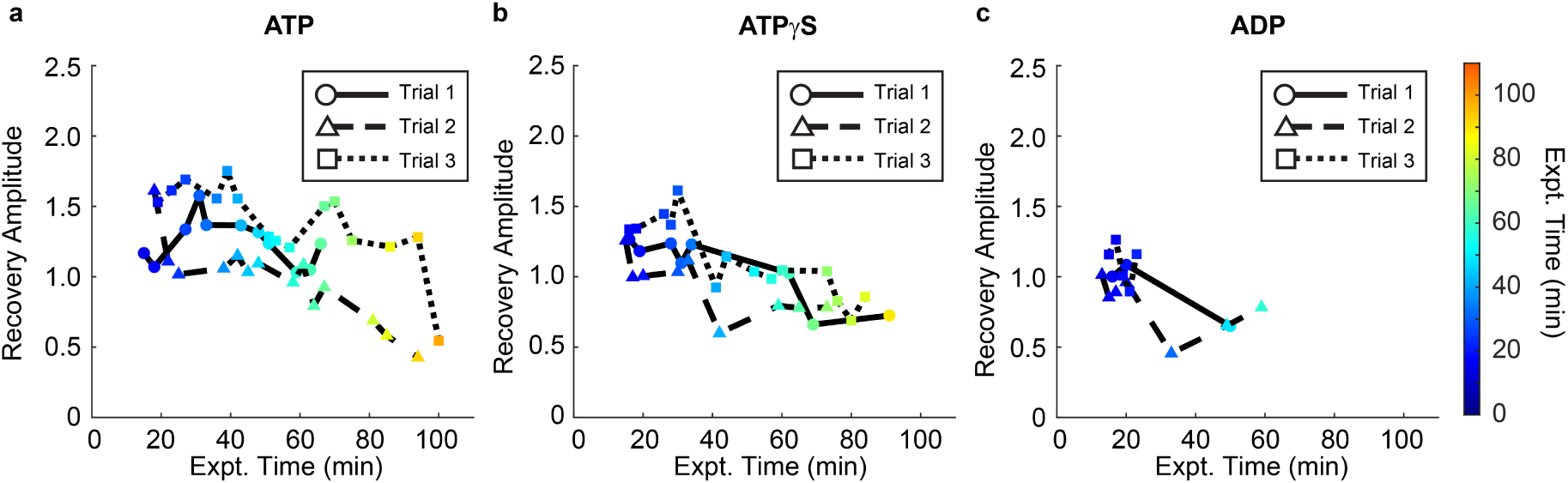
Hairpin RNA mobility within LAF-1 condensates decreases over time and depends on helicase activity. a-c) FRAP recovery amplitudes from the FRAP series of 4xMS2 hairpin RNA in LAF-1:RNA condensates shown in Figure 2. Condensates were formed in the presence of 1.6mM (a) ATP, (b) ATP*γ*S, or (c) ADP. A faster drop in recovery amplitude occurs as helicase activity is decreased. Recovery amplitudes greater than 1 are a result of RNA gradients in the condensed phase, with higher RNA concentrations at the center of the droplet.

**Figure S6:**
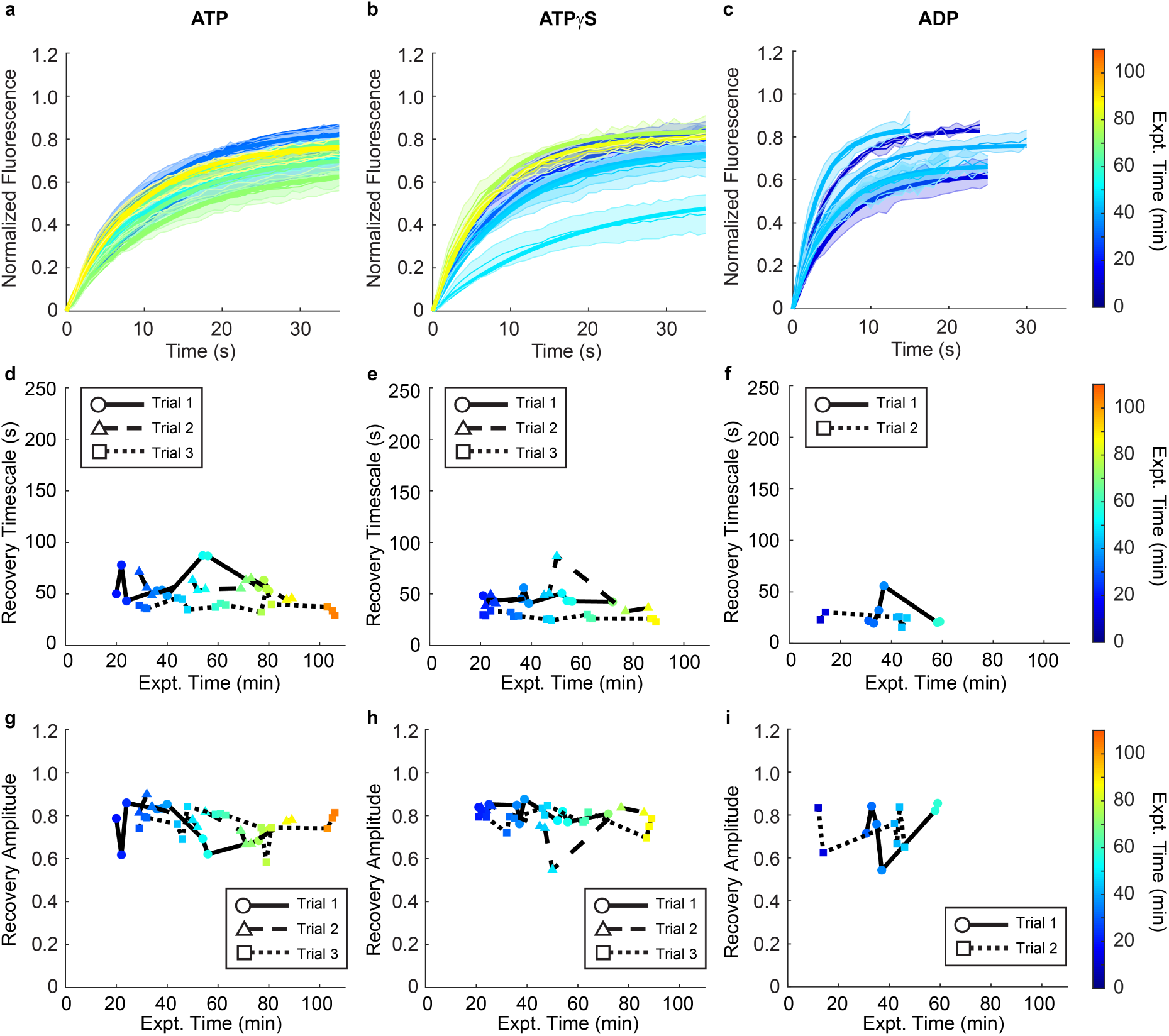
LAF-1 protein mobility in LAF-1:4xMS2 hairpin condensates does not change over time and is independent of helicase activity. a-c) FRAP timecourse series of LAF-1 protein in LAF-1:4xMS2 RNA condensates formed in the presence of 1.6mM (a) ATP, (b) ATP*γ*S, or (c) ADP. There is no difference in LAF-1 protein dynamics with different nucleotides present and LAF-1 protein dynamics do not change over time. d-f) Recovery timescale and (g-i) recovery amplitude as a function of the time at which the FRAP experiment was performed after system initialization for the indicated nucleotides. No difference in protein dynamics is seen between systems with different nucleotides, nor is there time-evolution of protein dynamics.

**Figure S7:**
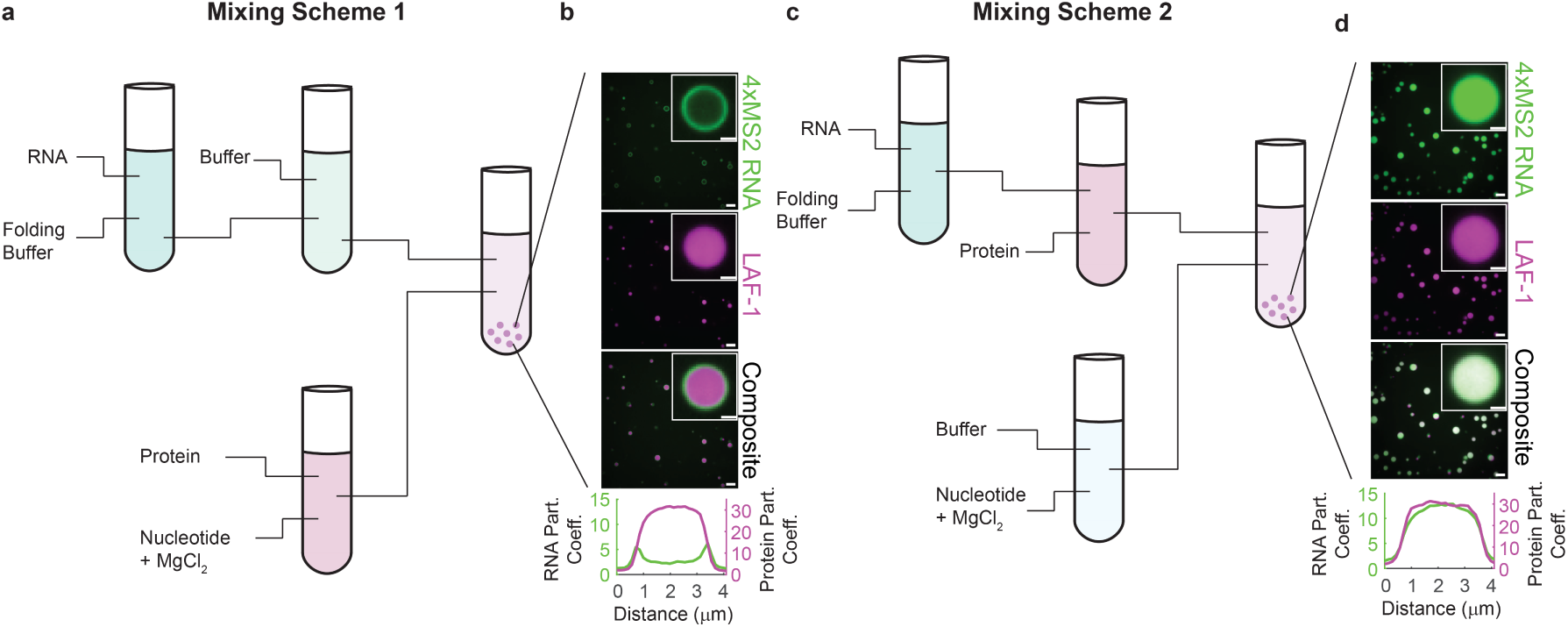
System mixing order affects droplet morphology. a) In the first reaction scheme, 4xMS2 RNA is annealed in buffer containing 10 mM Tris pH 7.4 and 10 mM KAc by heating to 95*^◦^*C and then cooled to room temperature. This RNA mixture is added to low salt buffer, and a mixture of LAF-1 protein in high salt buffer with nucleotide and magnesium are then added to initiate condensate formation. b) This reaction scheme results in core-shell structures of the droplets, where RNA accumulates at the LAF-1 condensate surface. This is believed to be due to RNA secondary structure formation prior to condensate formation. Scale bar of larger micrograph = 5 *µ*m, inset scale bar = 1 *µ*m. c) In the second reaction scheme, 4xMS2 RNA is annealed as described above, but first added to the LAF-1 protein in high salt buffer. This protein-RNA mixture is then added to low salt buffer containing ATP and magnesium to initiate condensate formation. d) This second reaction scheme produces initially homogeneous LAF-1:RNA co-condensates. We believe this to be due to condensate formation and RNA inclusion in the condensed phase prior to stable RNA secondary structure formation. Scale bar of larger micrograph = 5 *µ*m, inset scale bar = 1 *µ*m.

**Figure S8:**
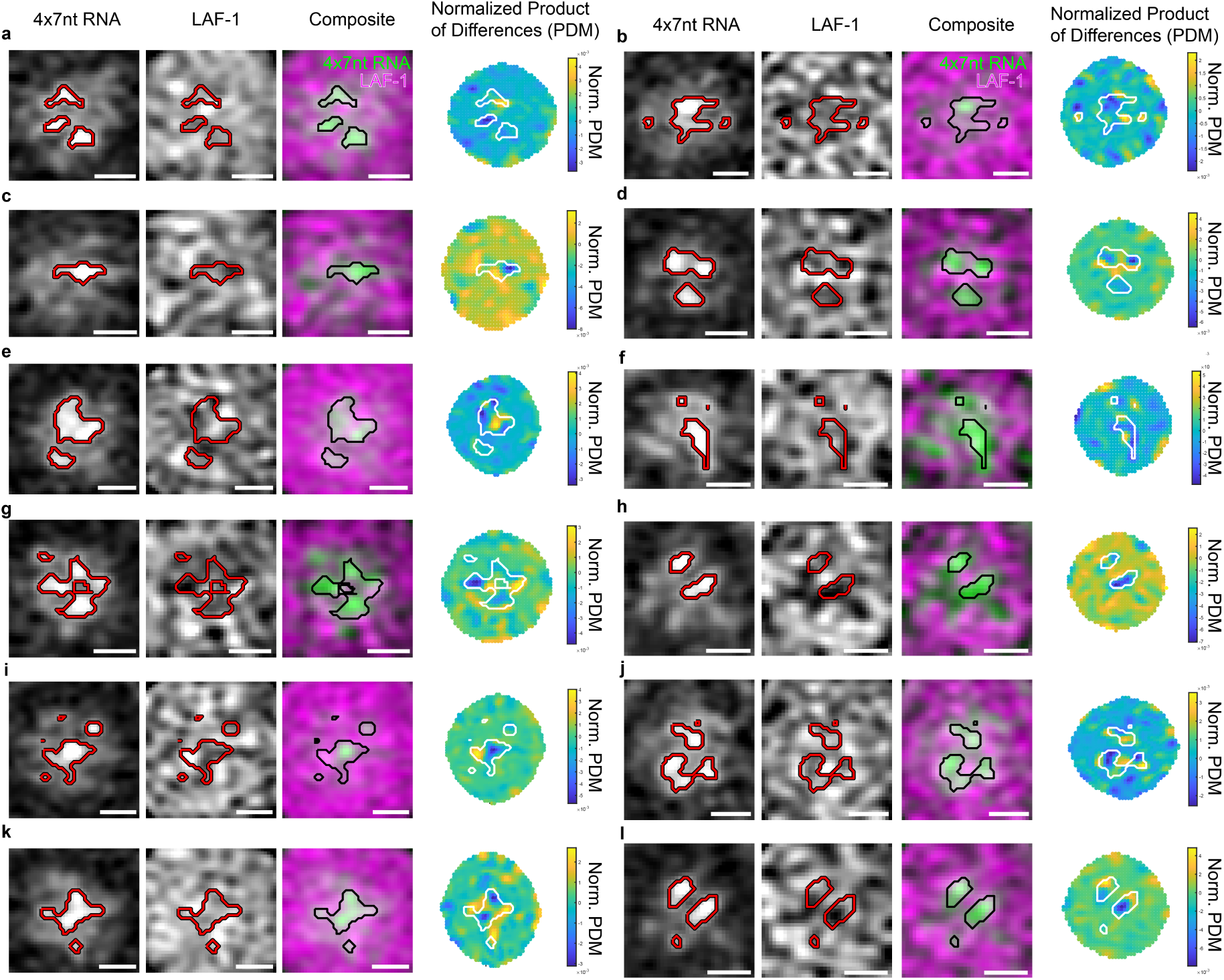
Normalized product of differences from the mean (PDM) analysis of different condensates. Additional examples of RNA punctate structures excluding LAF-1 protein from separate condensates. A more negative PDM value (more blue) indicates stronger anticorrelation between protein and RNA channels. Scale bar = 0.51 *µ*m. Related to Figure 3.

**Figure S9:**
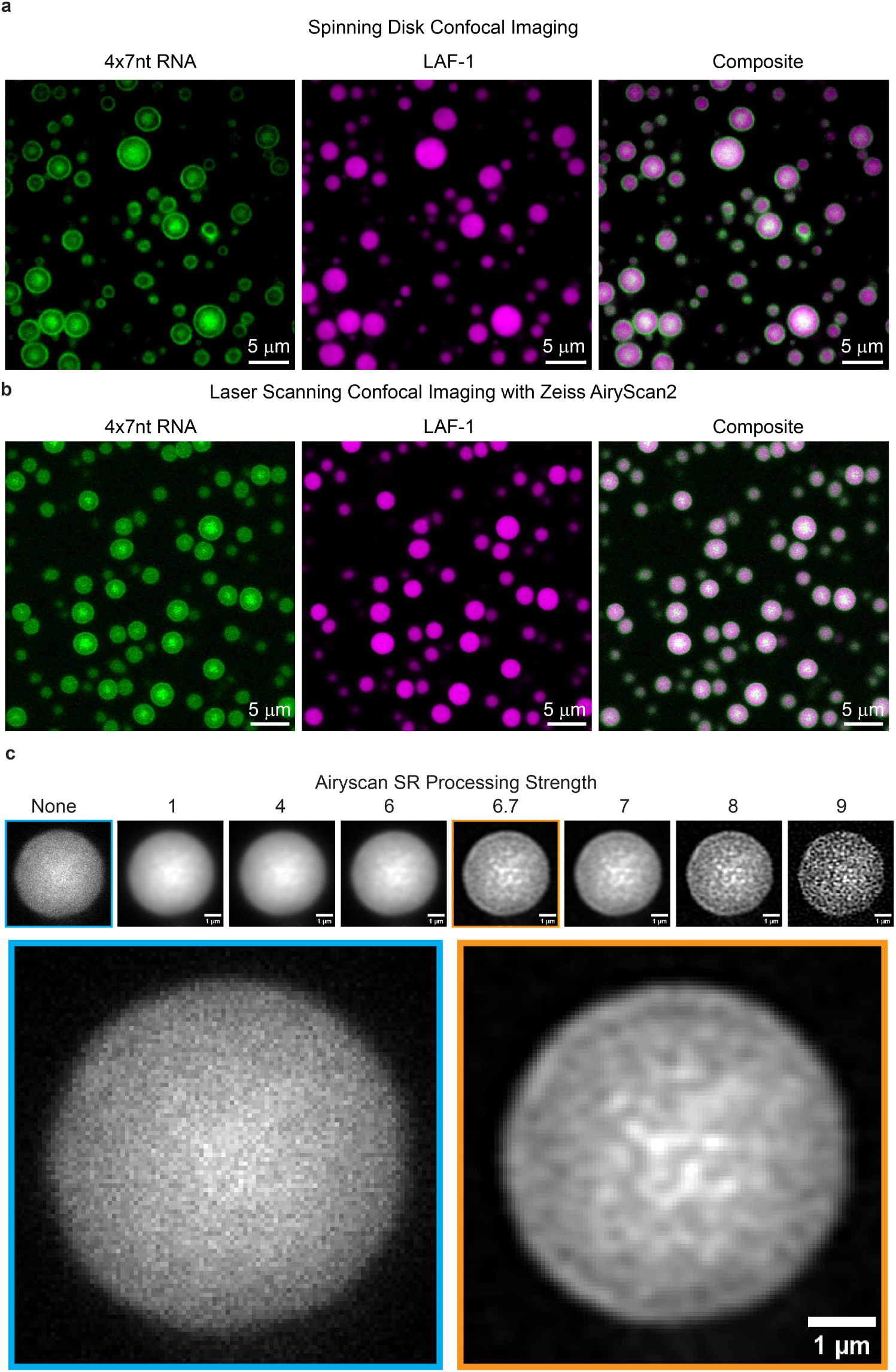
A coarse RNA structure is visible at the interior of LAF-1:4xMS2 RNA condensates after 2 hours. a) Imaging performed on a spinning disk confocal microscope shows coarse RNA structures at the core of LAF-1:4xMS2 condensates but with low spatial resolution. Scale bar = 2*µ*m. b) Imaging with a laser scanning confocal equipped with an Airyscan 2 detector increases spatial resolution of condensate imaging and shows RNA puncta at the core of LAF-1-RNA condensates. c) AiryScan processing strength changes the clarity of the spatial structure at the center of the condensate without introducing artificial spatial structure. Left to right shows the effect of increasing Airyscan processing strength. Shown below are larger images of the no Airyscan processing version (Left) and the image processed with the automatically detected Airyscan processing strength of 6.7 (Right).

**Figure S10:**
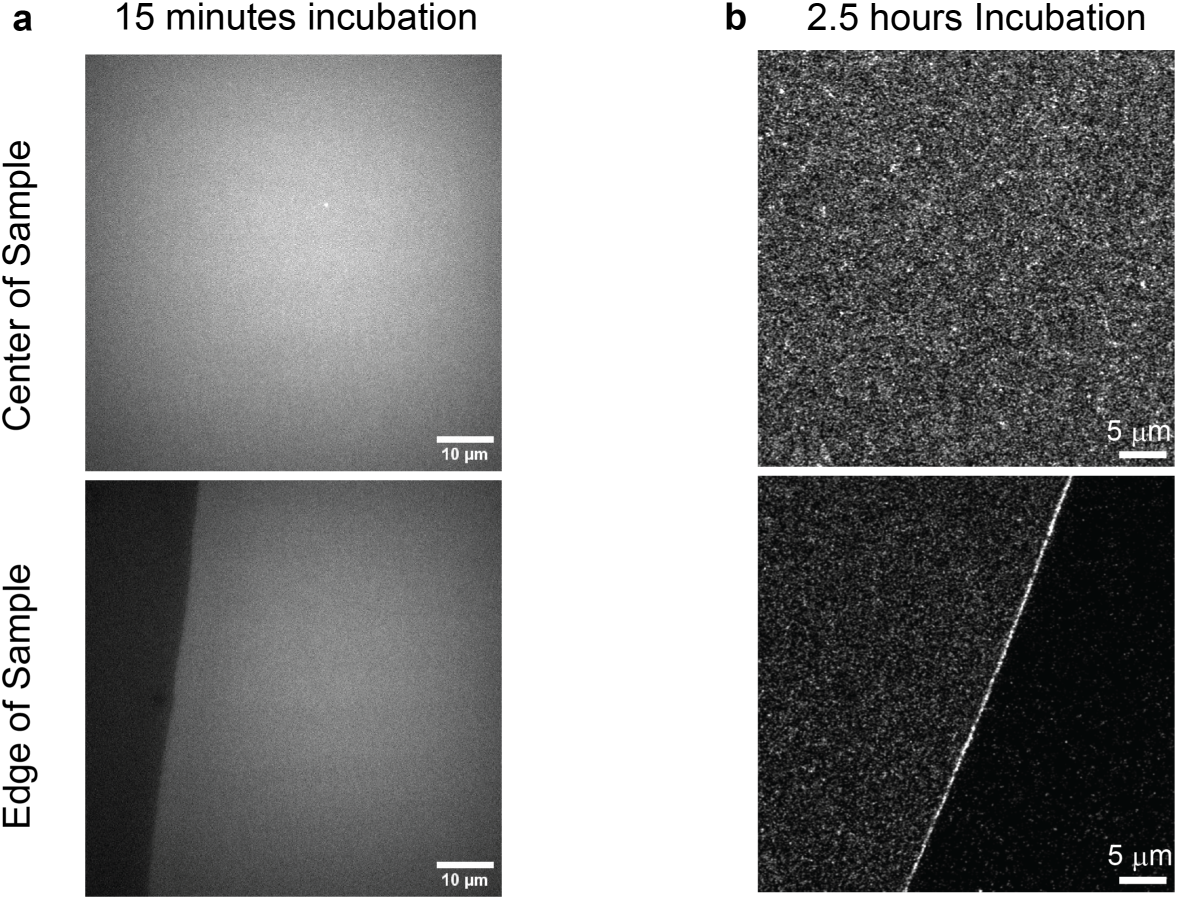
The 4xMS2 construct does not undergo aggregation or phase separation independent of the LAF-1 condensed phase. Under identical assay conditions to the condensate timecourses, but in the absence of LAF-1, 4xMS2 condensates do not form either initially (a) or after 2.5 hours of incubation (b). Micrographs in (a) were taken on a spinning disk confocal while the micrographs in (b) are from a Zeiss LSM980 with an Airyscan 2 detector. Scale bars correspond to: (a) 10 *µ*m, (b) are 5 *µ*m.

**Figure S11:**
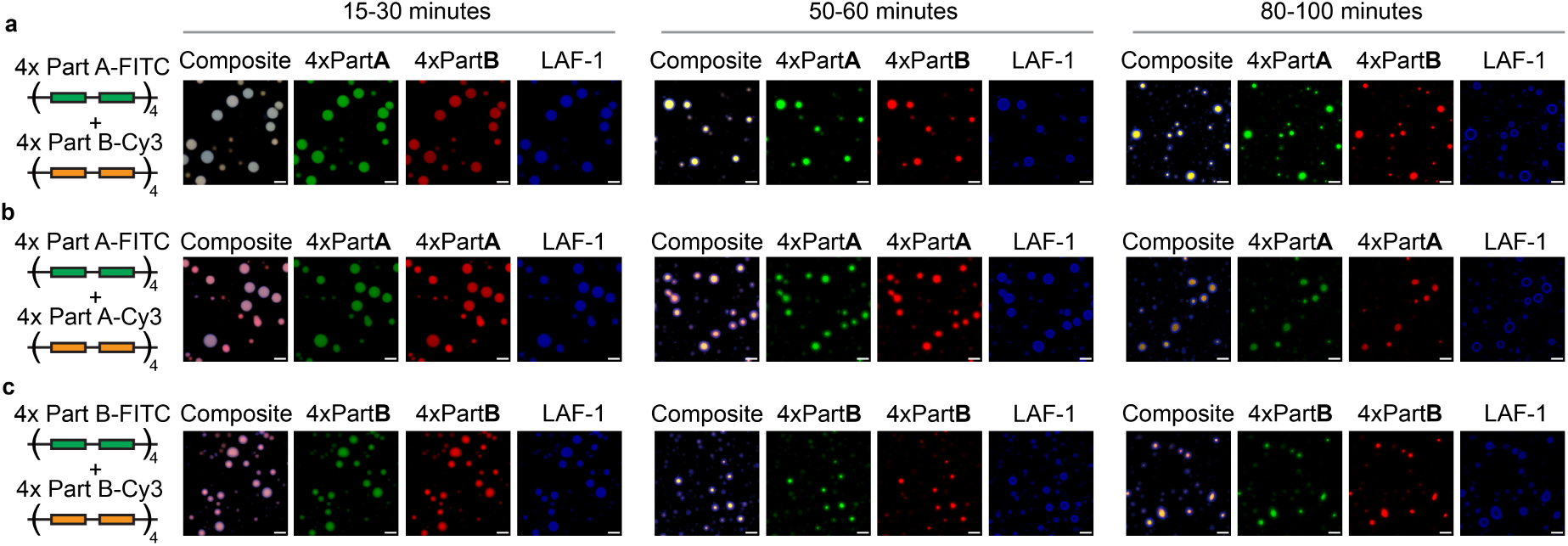
The second RNA phase transition does not depend on the specific base-pairing of the MS2 region. Separation between RNA-rich and protein rich phases is observed when both Part A and Part B of the MS2 hairpin sequence are present (a) but also when only tandem repeat of the Part A is present (b) or when tandem repeats of Part B are present (c). Though the condensates begin as homogeneous protein-RNA droplets, separation between protein- and RNA-rich phases occurs over 50-90 minutes in all cases. This suggests the full MS2 hairpin on a single RNA strand is helping to prevent coalescence of RNA within the condensed phase. Green channel indicates RNA labelled with fluorescein while the Red channel indicates RNA labelled with Cy3. Scale bar is equal to 5 *µ*m in all panels.

**Figure S12:**
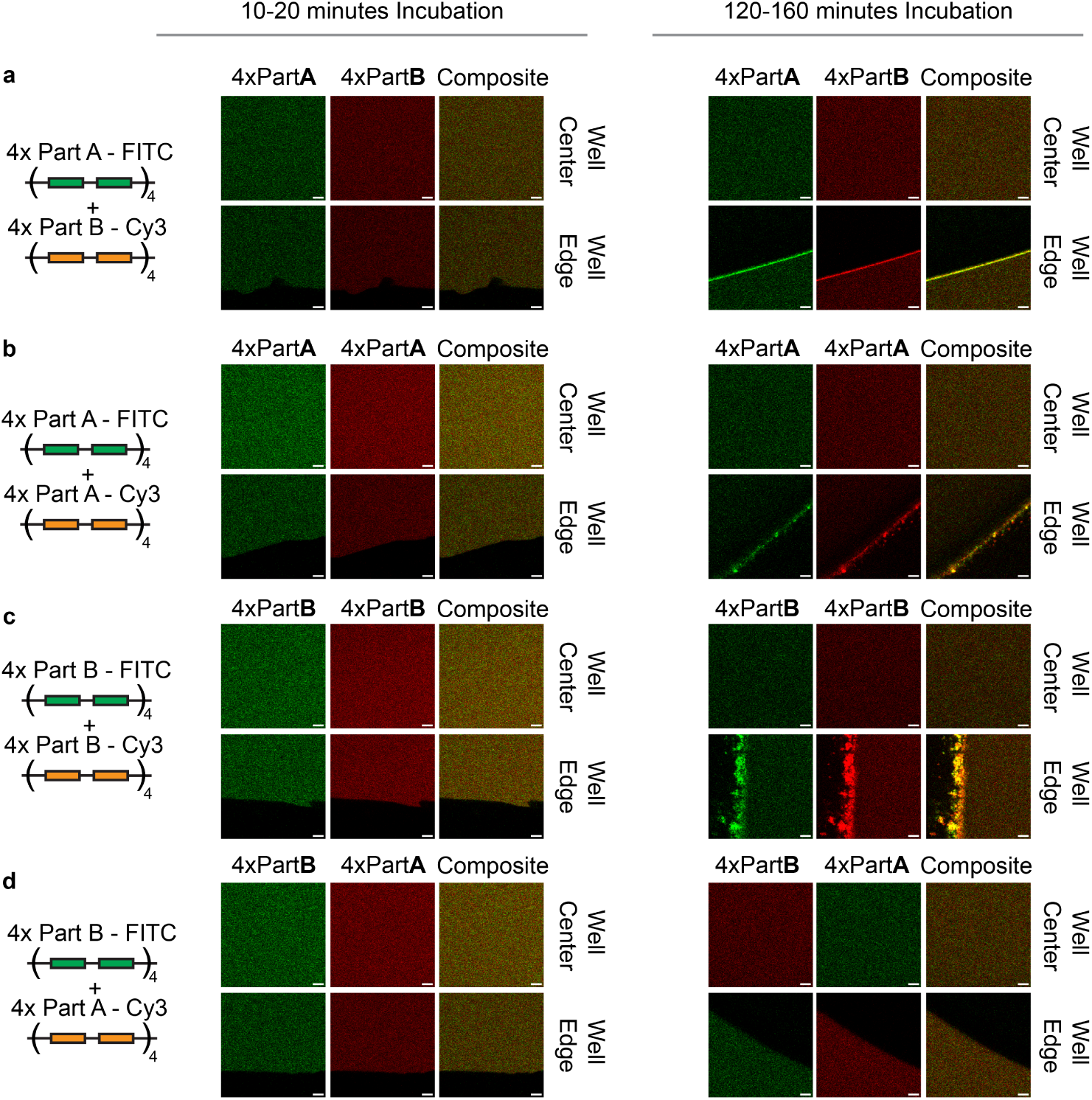
The different combinations of MS2 single-stranded sequences do not undergo phase separation or aggregation in the absence of LAF-1. Combining tandem repeats of Part A and Part B of the MS2 hairpin sequence (a), only Part A of the MS2 hairpin sequence (b), only part B of the MS2 hairpin sequence (c), or part A and part B with different fluorescent tags (d), do not form aggregates in the absence of LAF-1 either initially or over 2 hours. Green or left hand channel indicates RNA labelled with fluorescein while the Red or center channel indicates RNA labelled with Cy3. Some aggregation can be seen at the well edge after 2 hours but is not seen in the bulk. Scale bar is equal to 5 *µ*m in all panels.

## References

[1] Anthony A. Hyman, Christoph A. Weber, and Frank Jülicher. Liquid-liquid phase separation in biology. Annual Review of Cell and Developmental Biology, 30 (1): 39–58, 2014. doi: 10.1146/annurev-cellbio-100913-013325. URL 10.1146/annurev-cellbio-100913-013325. PMID: 25288112.

[2] Salman F. Banani, Hyun O. Lee, Anthony A. Hyman, and Michael K. Rosen. Biomolecular condensates: organizers of cellular biochemistry. Nature Reviews Molecular Cell Biology, 18 (5): 285–298, May 2017. ISSN 1471-0080. doi: 10.1038/nrm.2017.7. URL 10.1038/nrm.2017.7.

[3] Simon Alberti and Anthony A. Hyman. Biomolecular condensates at the nexus of cellular stress, protein aggregation disease and ageing. Nature Reviews Molecular Cell Biology, 22 (3): 196–213, Mar 2021. ISSN 1471-0080. doi: 10.1038/s41580-020-00326-6. URL -10.1038/s41580-020-00326-6.

[4] Louise Jawerth, Elisabeth Fischer-Friedrich, Suropriya Saha, Jie Wang, Titus Franzmann, Xiaojie Zhang, Jenny Sachweh, Martine Ruer, Mahdiye Ijavi, Shambaditya Saha, Julia Mahamid, Anthony A. Hyman, and Frank Jülicher. Protein condensates as aging Maxwell fluids. Science, 370 (6522): 1317–1323, 2020. doi: 10.1126/science.aaw4951. URL https://www.science.org/doi/abs/10.1126/science.aaw4951.

[5] Adiran Garaizar, Jorge R. Espinosa, Jerelle A. Joseph, Georg Krainer, Yi Shen, Tuomas P.J. Knowles, and Rosana Collepardo-Guevara. Aging can transform single-component protein condensates into multiphase architectures. Proceedings of the National Academy of Sciences, 119 (26): e2119800119, 2022. doi: 10.1073/pnas.2119800119. URL https://www.pnas.org/doi/abs/10.1073/pnas.2119800119.

[6] Sayantan Chatterjee, Yelena Kan, Mateusz Brzezinski, Kaloian Koynov, Roshan Mammen Regy, Anastasia C. Murthy, Kathleen A. Burke, Jasper J. Michels, Jeetain Mittal, Nicolas L. Fawzi, and Sapun H. Parekh. Reversible Kinetic Trapping of FUS Biomolecular Condensates. Advanced Science, 9 (4): 2104247, 2022. doi: 10.1002/advs.202104247. URL https://onlinelibrary.wiley.com/doi/abs/10.1002/-advs.202104247.

[7] Aravind Chandrasekaran, Kristin Graham, Jeanne C. Stachowiak, and Padmini Rangamani. Kinetic trapping organizes actin filaments within liquid-like protein droplets. Nature Communications, 15 (1): 3139, Apr 2024. ISSN 2041-1723. doi: 10.1038/s41467-024-46726-6. URL 10.1038/s41467-024-46726-6.

[8] Jennifer T Wang, Jarrett Smith, Bi-Chang Chen, Helen Schmidt, Dominique Rasoloson, Alexandre Paix, Bramwell G Lambrus, Deepika Calidas, Eric Betzig, and Geraldine Seydoux. Regulation of RNA granule dynamics by phosphorylation of serine-rich, intrinsically disordered proteins in *C. elegans*. eLife, 3: e04591, dec 2014. ISSN 2050-084X. doi: 10.7554/eLife.04591. URL 10.7554/eLife.04591.

[9] Andrew W. Folkmann, Andrea Putnam, Chiu Fan Lee, and Geraldine Seydoux. Regulation of biomolecular condensates by interfacial protein clusters. Science, 373 (6560): 1218–1224, 2021. doi: 10.1126/science.abg7071. URL https://www.science.org/doi/abs/-10.1126/science.abg7071.

[10] Yang Eric Guo, John C. Manteiga, Jonathan E. Henninger, Benjamin R. Sabari, Alessandra Dall’Agnese, Nancy M. Hannett, Jan-Hendrik Spille, Lena K. Afeyan, Alicia V. Zamudio, Krishna Shrinivas, Brian J. Abraham, Ann Boija, Tim-Michael Decker, Jenna K. Rimel, Charli B. Fant, Tong Ihn Lee, Ibrahim I. Cisse, Phillip A. Sharp, Dylan J. Taatjes, and Richard A. Young. Pol II phosphorylation regulates a switch between transcriptional and splicing condensates. Nature, 572 (7770): 543–548, Aug 2019. ISSN 1476-4687. doi: 10.1038/s41586-019-1464-0. URL 10.1038/s41586-019-1464-0.

[11] Jonathan E. Henninger, Ozgur Oksuz, Krishna Shrinivas, Ido Sagi, Gary LeRoy, Ming M. Zheng, J. Owen Andrews, Alicia V. Zamudio, Charalampos Lazaris, Nancy M. Hannett, Tong Ihn Lee, Phillip A. Sharp, Ibrahim I. Cissé, Arup K. Chakraborty, and Richard A. Young. RNA-Mediated Feedback Control of Transcriptional Condensates. Cell, 184 (1): 207–225.e24, Jan 2021. ISSN 0092-8674. doi: 10.1016/j.cell.2020.11.030. URL 10.1016/-j.cell.2020.11.030.

[12] Christopher Frederick Mugler, Maria Hondele, Stephanie Heinrich, Ruchika Sachdev, Pascal Vallotton, Adriana Y Koek, Leon Y Chan, and Karsten Weis. ATPase activity of the DEAD-box protein Dhh1 controls processing body formation. eLife, 5: e18746, 10 2016. ISSN 2050-084X. doi: 10.7554/eLife.18746. URL 10.7554/eLife.18746.

[13] Maria Hondele, Ruchika Sachdev, Stephanie Heinrich, Juan Wang, Pascal Vallotton, Beatriz M. A. Fontoura, and Karsten Weis. DEAD-box ATPases are global regulators of phase-separated organelles. Nature, 573 (7772): 144–148, 2019. ISSN 1476-4687. doi: 10.1038/s41586-019-1502-y. URL 10.1038/s41586-019-1502-y.

[14] Jan Kirschbaum and David Zwicker. Controlling biomolecular condensates via chemical reactions. Journal of The Royal Society Interface, 18 (179): 20210255, 2021. doi: 10.1098/rsif.2021.0255. URL https://royalsocietypublishing.org/doi/abs/10.1098/rsif.2021.0255.

[15] Christoph A Weber, Chiu Fan Lee, and Frank Jülicher. Droplet ripening in concentration gradients. New Journal of Physics, 19 (5): 053021, may 2017. doi: 10.1088/1367-2630/aa6b84. URL 10.1088/1367-2630/aa6b84.

[16] Clifford P. Brangwynne, Christian R. Eckmann, David S. Courson, Agata Rybarska, Carsten Hoege, Jöbin Gharakhani, Frank Jülicher, and Anthony A. Hyman. Germline P granules are liquid droplets that localize by controlled dissolution/condensation. Science, 324 (5935): 1729–1732, 2009. ISSN 0036-8075. doi: 10.1126/science.1172046. URL https://science.sciencemag.org/content/324/5935/1729.

[17] Jonathan Bauermann, Giacomo Bartolucci, Job Boekhoven, Christoph A. Weber, and Frank Jülicher. Formation of liquid shells in active droplet systems, 2023.

[18] Alexander M. Bergmann, Jonathan Bauermann, Giacomo Bartolucci, Carsten Donau, Michele Stasi, Anna-Lena Holtmannspötter, Frank Jülicher, Christoph A. Weber, and Job Boekhoven. Liquid spherical shells are a non-equilibrium steady state. bioRxiv, 2023. doi: 10.1101/2023.01.31.526480. URL https://www.biorxiv.org/content/early/2023/02/03/-2023.01.31.526480.

[19] David Zwicker, Rabea Seyboldt, Christoph A. Weber, Anthony A. Hyman, and Frank Jülicher. Growth and division of active droplets provides a model for protocells. Nature Physics, 13 (4): 408–413, Apr 2017. ISSN 1745-2481. doi: 10.1038/nphys3984. URL 10.1038/nphys3984.

[20] Miriam Linsenmeier, Maria Hondele, Fulvio Grigolato, Eleonora Secchi, Karsten Weis, and Paolo Arosio. Dynamic arrest and aging of biomolecular condensates are modulated by low-complexity domains, RNA and biochemical activity. Nature Communications, 13 (1): 3030, May 2022. ISSN 2041-1723. doi: 10.1038/s41467-022-30521-2. URL 10.1038/-s41467-022-30521-2.

[21] Inga Jarmoskaite and Rick Russell. DEAD-box proteins as RNA helicases and chaperones. Wiley Interdisciplinary Reviews: RNA, 2 (1): 135–152, 2011. doi: 10.1002/wrna.50. URL https://onlinelibrary.wiley.com/doi/abs/10.1002/wrna.50.

[22] Patrick Linder and Eckhard Jankowsky. From unwinding to clamping – the DEAD box RNA helicase family. Nature Reviews Molecular Cell Biology, 12 (8): 505–516, 2011. ISSN 1471-0080. doi: 10.1038/nrm3154. URL 10.1038/nrm3154.

[23] Devin Tauber, Gabriel Tauber, Anthony Khong, Briana Van Treeck, Jerry Pelletier, and Roy Parker. Modulation of rna condensation by the dead-box protein eif4a. Cell, 180 (3): 411–426.e16, Feb 2020. ISSN 0092-8674. doi: 10.1016/j.cell.2019.12.031. URL 10.1016/j.cell.2019.12.031.

[24] Ruchika Sachdev, Maria Hondele, Miriam Linsenmeier, Pascal Vallotton, Christopher F Mugler, Paolo Arosio, and Karsten Weis. Pat1 promotes processing body assembly by enhancing the phase separation of the DEAD-box ATPase Dhh1 and RNA. eLife, 8: e41415, 1 2019. ISSN 2050-084X. doi: 10.7554/eLife.41415. URL 10.7554/eLife.41415.

[25] Sebastian Coupe and Nikta Fakhri. ATP-induced cross-linking of a biomolecular condensate. Biophysical Journal, 2023/08/10 XXXX. ISSN 0006-3495. doi: 10.1016/j.bpj.2023.07.013. URL 10.1016/j.bpj.2023.07.013.

[26] Kyle Begovich and James E. Wilhelm. An In Vitro Assembly System Identifies Roles for RNA Nucleation and ATP in Yeast Stress Granule Formation. Molecular Cell, 79 (6): 991–1007.e4, Sep 2020. ISSN 1097-2765. doi: 10.1016/j.molcel.2020.07.017. URL 10.1016/j.molcel.2020.07.017.

[27] Christine Roden and Amy S. Gladfelter. RNA contributions to the form and function of biomolecular condensates. Nature Reviews Molecular Cell Biology, 22 (3): 183–195, Mar 2021. ISSN 1471-0080. doi: 10.1038/s41580-020-0264-6. URL 10.1038/s41580-020-0264-6.

[28] Shana Elbaum-Garfinkle, Younghoon Kim, Krzysztof Szczepaniak, Carlos Chih-Hsiung Chen, Christian R. Eckmann, Sua Myong, and Clifford P. Brangwynne. The disordered P granule protein LAF-1 drives phase separation into droplets with tunable viscosity and dynamics. Proceedings of the National Academy of Sciences, 112 (23): 7189–7194, 2015. ISSN 0027-8424. doi: 10.1073/pnas.1504822112. URL https://www.pnas.org/content/112/23/7189.

[29] Ming-Tzo Wei, Shana Elbaum-Garfinkle, Alex S. Holehouse, Carlos Chih-Hsiung Chen, Marina Feric, Craig B. Arnold, Rodney D. Priestley, Rohit V. Pappu, and Clifford P. Brangwynne. Phase behaviour of disordered proteins underlying low density and high permeability of liquid organelles. Nature Chemistry, 9: 1118, 6 2017. URL 10.1038/nchem.2803. Article.

[30] Shovamayee Maharana, Jie Wang, Dimitrios K. Papadopoulos, Doris Richter, Andrey Pozniakovsky, Ina Poser, Marc Bickle, Sandra Rizk, Jordina Guillén-Boixet, Titus M. Franzmann, Marcus Jahnel, Lara Marrone, Young-Tae Chang, Jared Sterneckert, Pavel Tomancak, Anthony A. Hyman, and Simon Alberti. RNA buffers the phase separation behavior of prion-like RNA binding proteins. Science, 360 (6391): 918–921, 2018. doi: 10.1126/science.aar7366. URL https://www.science.org/doi/abs/10.1126/science.aar7366.

[31] Ankur Jain and Ronald D. Vale. RNA phase transitions in repeat expansion disorders. Nature, 546: 243, 5 2017. URL 10.1038/nature22386. Article.

[32] Sumit Majumder, Sebastian Coupe, Nikta Fakhri, and Ankur Jain. Sequence programmable nucleic acid coacervates. bioRxiv, 2024. doi: 10.1101/2024.07.22.604687. URL https://www.biorxiv.org/content/early/2024/07/23/2024.07.22.604687.

[33] Saehyun Choi, McCauley O. Meyer, Philip C. Bevilacqua, and Christine D. Keating. Phase-specific RNA accumulation and duplex thermodynamics in multiphase coacervate models for membraneless organelles. Nature Chemistry, 14 (10): 1110–1117, Oct 2022. ISSN 1755-4349. doi: 10.1038/s41557-022-00980-7. URL 10.1038/s41557-022-00980-7.

[34] Raghav R Poudyal, Jacob P Sieg, Bede Portz, Christine D Keating, and Philip C Bevilacqua. RNA sequence and structure control assembly and function of RNA condensates. RNA, 27 (12): 1589–1601, December 2021.

[35] Avinash Patel, Liliana Malinovska, Shambaditya Saha, Jie Wang, Simon Alberti, Yamuna Krishnan, and Anthony A. Hyman. ATP as a biological hydrotrope. Science, 356 (6339): 753–756, 2017. ISSN 0036-8075. doi: 10.1126/science.aaf6846. URL https://science.sciencemag.org/content/356/6339/753.

[36] Younghoon Kim and Sua Myong. RNA remodeling activity of DEAD box proteins tuned by protein concentration, RNA length, and ATP. Molecular Cell, 63 (5): 865–876, Sep 2016. ISSN 1097-2765. doi: 10.1016/j.molcel.2016.07.010. URL 10.1016/-j.molcel.2016.07.010.

[37] Warren A. Kibbe. OligoCalc: an online oligonucleotide properties calculator. Nucleic Acids Research, 35 (suppl2): W43–W46, 07 2007. ISSN 0305-1048. doi: 10.1093/nar/gkm234. URL 10.1093/nar/gkm234.

[38] Tharun Selvam Mahendran, Gable M Wadsworth, Anurag Singh, and Priya R Banerjee. Biomolecular Condensates Can Enhance Pathological RNA Clustering. bioRxiv, 2024. doi: 10.1101/2024.06.11.598371. URL https://www.biorxiv.org/content/early/2024/06/13/-2024.06.11.598371.

[39] Ali R. Özes, Kateryna Feoktistova, Brian C. Avanzino, Enoch P. Baldwin, and Christopher S. Fraser. Real-time fluorescence assays to monitor duplex unwinding and ATPase activities of helicases. Nature Protocols, 9: 1645 EP –, Jun 2014. URL 10.1038/-nprot.2014.112.

[40] Edouard Bertrand, Pascal Chartrand, Matthias Schaefer, Shailesh M. Shenoy, Robert H. Singer, and Roy M. Long. Localization of ASH1 mRNA Particles in Living Yeast. Molecular Cell, 2 (4): 437–445, 1998. ISSN 1097-2765. doi: 10.1016/S1097-2765(00)80143-4. URL https://www.sciencedirect.com/science/article/pii/S1097276500801434.

[41] Nicole O. Taylor, Ming-Tzo Wei, Howard A. Stone, and Clifford P. Brangwynne. Quantifying dynamics in phase-separated condensates using fluorescence recovery after photobleaching. Biophysical Journal, 117 (7): 1285–1300, 2019. ISSN 0006-3495. doi: 10.1016/j.bpj.2019.08.030. URL https://www.sciencedirect.com/science/article/-pii/S0006349519307477.

## References

1. Özes, A. R., Feoktistova, K., Avanzino, B. C., Baldwin, E. P. & Fraser, C. S. Real-time fluorescence assays to monitor duplex unwinding and ATPase activities of helicases. Nature Protocols 9, 1645 EP -. 10.1038/nprot.2014.112 (June 2014).

2. Linder, P. & Jankowsky, E. From unwinding to clamping – the DEAD box RNA helicase family. Nature Reviews Molecular Cell Biology 12, 505–516. issn: 1471-0080. 10.1038/nrm3154 (2011).

